# Thymidine starvation promotes c-di-AMP dependent inflammation during infection

**DOI:** 10.1101/2020.08.21.261750

**Authors:** Qing Tang, Mimi R. Precit, Maureen K. Thomason, Fariha Ahmed-Qadri, Adelle P. McFarland, Daniel J. Wolter, Lucas R. Hoffman, Joshua J. Woodward

## Abstract

Antibiotics remain one of the most effective methods for controlling bacterial infection. However, the diverse impacts of antimicrobials on bacterial physiology and host immunity remain unclear. A comprehensive antibiotic screen revealed that disruption of thymidine synthesis in Firmicutes with anti-folate antibiotics promoted elevated levels of the bacterial second messenger cyclic di-AMP, and consequently induced host STING activation during infection. Extensive exposure to antibiotics targeting folate synthesis drives the emergence of thymidine-dependent *Staphylococcus aureus* SCVs (TD-SCVs). Respiratory infections with TD-SCVs are common among children with cystic fibrosis and are associated with worse clinical outcomes, although the underlying pathophysiological mechanisms remain to be defined. Our study reveals that TD-SCV isolates exhibited excessive c-di-AMP production and STING activation in a thymidine-dependent manner. Murine lung infection with TD-SCVs revealed STING-dependent elevation of proinflammatory cytokines, leading to higher airway neutrophil infiltration and activation comparing to normal colony *S. aureus* and hemin-dependent SCV. Our results suggest the elevated inflammatory capacity of TD-SCVs contribute to their pathogenesis and revealed a new aspect of STING signaling in the airway by characterizing its role in neutrophil recruitment.

## Introduction

The discovery of antibiotics over one hundred years ago marked one of the most significant advancements in human medical interventions of the 20th century, providing a cure for many otherwise fatal bacterial infections and profoundly impacting human morbidity and mortality. Since the initial discovery of penicillin, we have faced countless challenges in antibiotic utilization, most notably the acquisition of antibiotic resistance. The development of new antimicrobial classes characterized by their disruption of distinct aspects of bacterial physiology continues to this day. *In vitro*, these compounds inhibit microbial growth or directly kill microbes, depending upon their mechanism of action. In the *in vivo* context, it has long been appreciated that these therapeutic agents can also directly affect the host immune system, in many cases augmenting microbial growth inhibition and in some instances leading to pathological inflammation, most notably toxic shock^1^. Despite decades of study, the complex relationship between antimicrobials, bacterial physiology, and host immunity remains incompletely understood.

Cyclic dinucleotides (CDNs) of bacterial origin play key roles in mediating host immune responses to infection. Earlier studies identified secretion of c-di-AMP by *Listeria monocytogenes* during infection as a conserved microbial signature for innate immune detection of several pathogens^2^. The host protein STING is now recognized as the receptor for detecting bacterial cyclic dinucleotides (c-di-AMP, c-di-GMP and 3′3′-cGAMP), in addition to eukaryotic 2′3′-cGAMP, which is synthesized by cGAS in response to host or pathogen DNA^3–7^. STING binding to cyclic di-nucleotides promotes IRF3 phosphorylation and translocation to the nucleus to mediate transcription of IFN-β and other coregulated genes^3^. Additionally, STING activation promotes NF-κB, MAP kinase, STAT6, NLRP3 inflammasome activation, apoptosis, and autophagy, independent of IFN-β^8–12^. Collectively, the CDN-STING signaling axis has emerged as a central mediator of host immunity to microbial infection in a variety of contexts^13–15^. However, aberrant STING activation has also been linked to the development of several autoimmune disorders affecting the brain, vasculature, and lungs^16–18^. It is currently unclear if STING activation is impacted by antimicrobial treatment or how such treatment impacts infection outcome.

Bacteria encode countless systems to ensure their survival in the face of varying essential nutrient availability and environmental conditions, and with disruption of central cellular processes. In addition to its role as a key mediator of inflammation, the second messenger c-di-AMP is now appreciated to play a pleiotropic role in many such survival responses, with direct impacts on cell wall synthesis, regulation of osmolyte levels, DNA damage repair, and notably resistance to cell wall-targeting antibiotics^19,20^. Accumulation of c-di-AMP by mutation of its phosphodiesterase GdpP mediates β-lactam resistance in clinical isolates of *Staphylococcus aureus*^21–23^. Additionally, other important mechanisms of antibiotic resistance have been documented in both clinical and laboratory settings. Horizontal transfer of antimicrobial resistance genes has garnered widespread attention^24^. However, in many organisms, including *S. aureus*, antibiotic therapy can drive the emergence of isolates harboring disruptions in genes involved in fundamental metabolic pathways. The resulting small colony variants (SCVs) exhibit slow *in vitro* growth, and *S. aureus* SCVs are associated with antibiotic-refractory, chronic infections, including osteomyelitis, endocarditis, wound infections, and lung infections in patients with the autosomal recessive disease cystic fibrosis (CF)^25,26^. As such, SCV infections are difficult to eradicate and can persist for many years. In some contexts, notably among adolescent CF patients, the presence of SCVs is associated with worse disease outcomes^27,28^. Despite our rapidly expanding understanding of c-di-AMP function in microbial physiology, the impact of this signaling molecule on the central processes disrupted by antibiotic exposure and those involved in antibiotic resistance has not been defined.

In this study, we conducted a screen to characterize the impacts of antibiotic exposure on STING-dependent inflammation and c-di-AMP production using the model intracellular pathogen *L. monocytogenes*. These studies showed that antibiotic disruption of thymidine metabolism among several human pathogens results in elevations in both c-di-AMP production and consequent STING-dependent inflammation. Additionally, *S. aureus* SCVs carrying mutations in the enzyme thymidine synthase, which are frequently detected in patients following antifolate antibiotic therapy, confer similar c-di-AMP and STING-dependent hyperinflammation and elevated inflammatory cell recruitment to the airway during infection of the lung. Collectively, these findings reveal an unappreciated link between antibiotic therapy, antifolate resistance, and host inflammation, providing initial insight into how these processes impact host outcomes during infection.

## Results

### Antibiotic stresses contribute to bacterial intracellular c-di-AMP accumulation

In many Firmicutes, c-di-AMP plays a central role in microbial stress responses and is essential for microbial viability in several growth conditions^19^. However, there remains a dearth of knowledge pertaining to the environmental cues that impact nucleotide dynamics in this phylum of bacteria. Given that antibiotics function as microbial stress agents that target central aspects of bacterial physiology, we hypothesized that bacterial exposure to these therapeutic agents could modulate nucleotide levels and illuminate c-di-AMP’s role in central microbial stress responses.

The intracellular pathogen *L. monocytogenes* secretes c-di-AMP, which activates STING and elicits IFN-β production^2^, providing an indirect readout of cyclic dinucleotide levels during infection. To explore if specific classes of antibiotics affect c-di-AMP production in this pathogen, we performed a screen by incubating *L. monocytogenes* in BIOLOG™ microplates (PM11C and PM12B) containing 48 different antibiotics at various concentrations. Bacteria were subsequently used to infect primary murine bone marrow derived macrophages (pBMDMs), and the resulting IFN-β production was measured by luciferase bioassay. *L. monocytogenes* treated with a variety of β-lactam antibiotics, which disrupt bacterial cell wall homeostasis, induced elevated IFN-β concentrations relative to the untreated control (Fig. 1a). We also observed an unexpected elevation of IFN-β following exposure to 2,4-Diamino-6,7-diisopropylpteridine, which targets folate synthesis (Fig. 1a-b). To validate and expand upon the observations from the BIOLOG™ antibiotic screen, *L. monocytogenes* was treated with other antibiotics that target folate synthesis, including sulfamethoxazole, trimethoprim, and SXT (trimethoprim/sulfamethoxazole combination in a 1:5 ratio, also known as Bactrim), as well as the cell wall-targeting antibiotic penicillin G. Bacteria were subsequently used to infect macrophages and IFN-β induction was determined by luciferase bioassay. Consistent with the initial screen, penicillin G induced elevated IFN-β production during *L. monocytogenes* infection. Additionally, antibiotics specifically targeting dihydrofolate reductase (DHFR), which converts dihydrofolate to tetrahydrofolate, induced significantly higher IFN-β production, whereas treatment with the dihydropteroate synthase (DHPS) inhibitor sulfamethoxazole, an enzyme involved in dihydrofolate synthesis, resulted in a comparable level of IFN-β relative to the untreated control (Fig. 1c). These observations reveal that antibiotics that specifically target tetrahydrofolate synthesis promote IFN-β production during infection by *L. monocytogenes*.

**Fig. 1:**
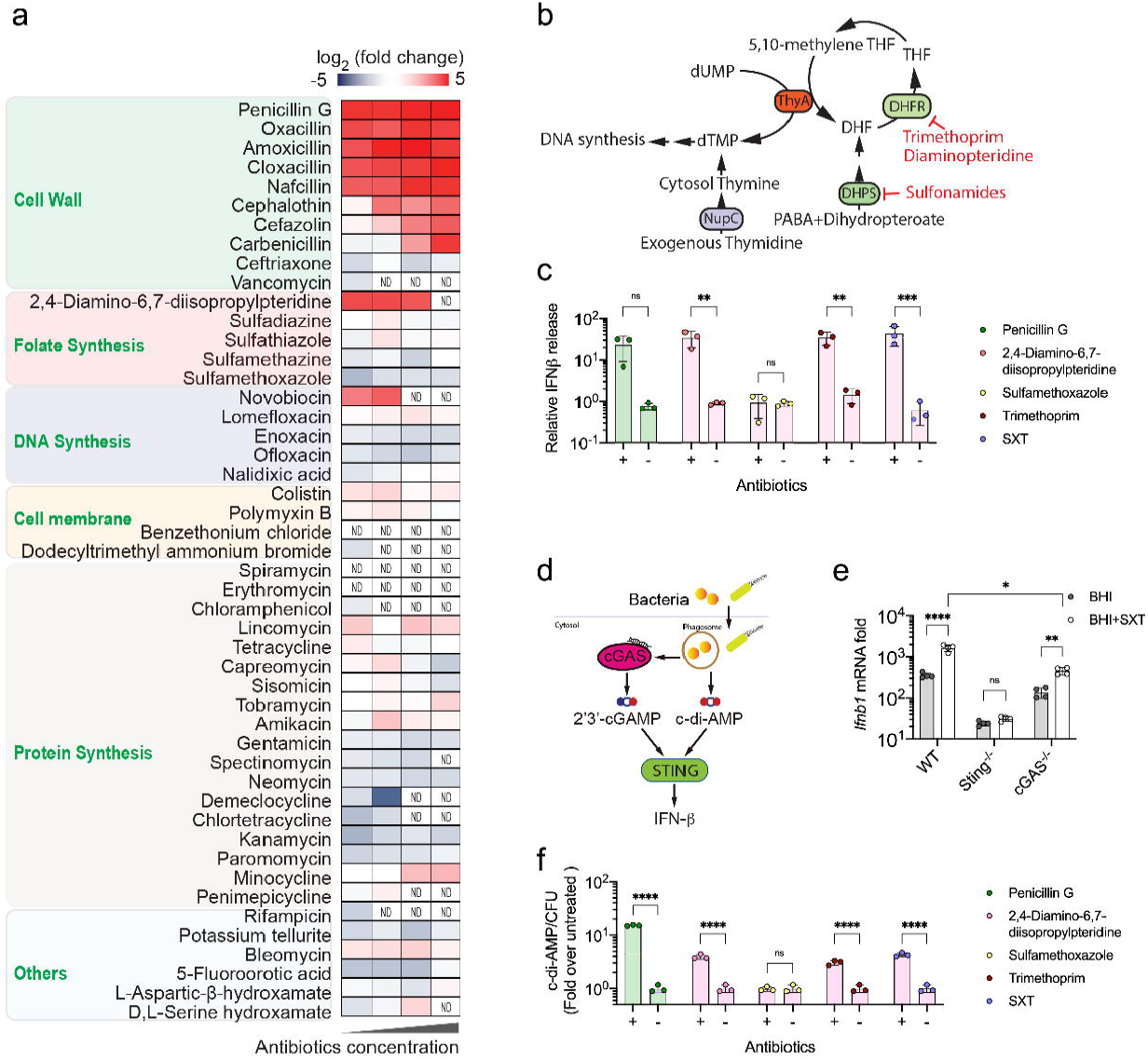
Antibiotic treatment promotes c-di-AMP production by bacteria. **a**, Heatmap of relative IFN-β production by pBMDMs infected with *L. monocytogenes* incubated in BioLog antibiotic plates (PM11C and PM12B). The IFN-β in macrophage supernatants was measured by luciferase bioassays using IFN-responsive ISRE-L929 cells. Values are the log_2_ ratio of IFN-β production relative to untreated *L. monocytogenes* control. Black labels indicate different antibiotics, and the green labels indicate the antibiotic class based on mechanism of action. ND indicates that no bacteria were recovered after antibiotic treatments and IFN-β was not determined. **b**, Schematic diagram of thymidine metabolism in bacteria. ThyA: Thymidylate synthase; NupC: Nucleoside permease; DHFR: Dihydrofolate reductase; DHPS: Dihydropteroate synthase. **c**, IFN-β production of pBMDMs infected with *L. monocytogenes* treated with indicated antibiotics relative to the untreated control at 6 hpi as measured by luciferase bioassays using IFN responsive ISRE-L929 cells. Logarithmic phase *L. monocytogenes* were incubated with either 15 μg/ml penicillin G, 200 μg/ml 2,4-Diamino-6,7-diisopropylpteridine, 1,200 μg/ml Sulfamethoxazole, 250 μg/ml Trimethoprim, or 50 μg/ml Bactrim for 6 hrs before macrophage phagocytosis. **d**, Schematic diagram of STING activation. Eukaryotic 2′3′-cGAMP, synthesized by cGAS that is triggered by host or pathogen-derived DNA, and bacterial cyclic dinucleotides can both active STING, resulting in IFN-β induction. **e**, *Ifnb1* mRNA expression in WT or *Sting* ^-/-^ pBMDMs infected with *L. monocytogenes* grown in BHI broth with or without 0.25 mg/ml of Bactrim supplementation at 4 hpi. **f**, Quantification of intracellular c-di-AMP production of *L. monocytogenes* treated with antibiotics as indicated in panel **c**) relative to the untreated control. For all panels, mean values of biological replicates are plotted, and error bars indicate ±SD, *P* values were calculated using Two-way ANOVA analysis. Asterisks indicate that differences are statistically significant (*, *P* < 0.05; **, *P* < 0.01; ***, *P* < 0.001, ****, *P* < 0.0001), and ns indicates no significant difference.

In the absence of antibiotics, IFN-β production during *L. monocytogenes* infection is primarily attributed to direct activation of STING by bacterial c-di-AMP. However, IFN-β production can also result from activation of cGAS by host or pathogen-derived DNA and subsequent 2′3′-cGAMP-mediated STING activation^6^ (Fig. 1d). To discern if IFN-β induction downstream of DHFR inhibition is due to direct bacterial STING antagonism or bacterial DNA release, WT, *Sting*^-/-^ and *cGas*^-/-^ pBMDMs were infected with *L. monocytogenes* cultured with or without a sub-inhibitory concentration of SXT. In both cases, IFN-β induction was greatly reduced in the absence of STING and there was an insignificant effect of SXT on cytokine production, confirming the role of this cyclic dinucleotide receptor regardless of antibiotic presence. Additionally, SXT-treated *L. monocytogenes* induced significantly higher *Ifbn1* transcription in both WT and *cGas*^-/-^ cells relative to the untreated control (Fig. 1e). Because antibiotic-induced *Ifbn1* expression was cGAS-independent, these observations support direct STING activation by *L. monocytogenes* following SXT exposure.

Given our observations that STING activation following SXT treatment is independent of cGAS sensing of DNA, and the known role of c-di-AMP as a direct STING activator during *L. monocytogenes* infection, our findings suggested that exposure to antibiotics that target DHFR results in increased CDN production. To confirm this assertion, we monitored c-di-AMP levels in *L. monocytogenes* treated with penicillin G, 2,4-diamino-6,7-diisopropylpteridine, sulfamethoxazole, trimethoprim and SXT. C-di-AMP levels directly mirrored the IFN-β induction observed following macrophage infection, whereby all of these antibiotics except for the DHPS inhibitor sulfamethoxazole, resulted in the significant elevation of nucleotide levels (Fig. 1f). Collectively, these observations reveal that antibiotics that target DHFR result in elevated c-di-AMP production by *L. monocytogenes*, which promote STING-dependent IFN-β production by infected host cells.

### Thymidine depletion promotes c-di-AMP production of Firmicutes

DHFR plays a key role in folate regeneration, which is required for several cellular biosynthetic pathways, including those for thymidine, amino acids, and purines^29^. The bactericidal activity of antibiotics targeting folate biosynthesis results from disrupting thymidine synthesis, which induces a lethal process known as thymineless death^30^. To determine if the effect of folate inhibition on c-di-AMP production was due to disrupted thymidine metabolism, we quantified bacterial c-di-AMP levels as well as macrophage IFN-β production during infection with *L. monocytogenes* grown under SXT stress in the presence or absence of exogenous thymidine. In the absence of thymidine, SXT induced a 15-fold elevation of c-di-AMP and IFN-β production, while thymidine supplementation complemented c-di-AMP and IFN-β levels induced by SXT exposure (Fig. 2a-b). These observations indicated that thymidine depletion following DHFR inhibition promotes c-di-AMP production. To further verify these results, we generated an in-frame deletion of the thymidylate synthase gene *thyA* in *L. monocytogenes* and compared *Ifnb1* induction during macrophage infection with this mutant to that by WT *L. monocytogenes*. Consistent with our observations with SXT, Δ*thyA* induced significantly higher *Ifnb1* expression relative to WT *L. monocytogenes* in a STING-dependent and cGAS-independent manner (Fig. 2c). These observations support the conclusion that altered thymidine biosynthesis, either through antibiotic-mediated inhibition of tetrahydrofolate regeneration or through disruption of *de novo* thymidine biosynthesis via genetic means, is responsible for the increased c-di-AMP production of *L. monocytogenes*. Whereas tetrahydrofolate is an essential cofactor for ThyA in all bacteria^31^, the di-adenylate cyclase (DacA) which synthesizes c-di-AMP is essential for Firmicutes but absent in most Proteobacteria^19^. To explore the generality of our observations linking thymidine metabolism and IFN-β production during infection, four common human pathogens were grown under sub-inhibitory concentrations of SXT in the presence or absence of surplus exogenous thymidine, and induction of *Ifnb1* transcription was assessed by qRT-PCR following macrophage infection. SXT treatment induced *Ifnb1* expression in a thymidine-dependent manner in the c-di-AMP-producing Firmicutes *Enterococcus faecalis* (Fig. 2d), *S. aureus* Newman (Fig. 2e), and *S. aureus* JE2 (Fig. 2f). However, the c-di-AMP deficient Proteobacteria *Salmonella enterica* serovar Typhimurium (Fig. 2g) and *Francisella novicida* (Fig. 2h) either induced comparable *Ifnb1* in all conditions or induced decreased *Ifnb1* after SXT treatment regardless of thymidine supplementation. These results provide evidence that disrupting thymidine metabolism increases c-di-AMP production in diverse c-di-AMP producing Firmicutes, but not among Proteobacteria that do not produce this second messenger.

**Fig. 2:**
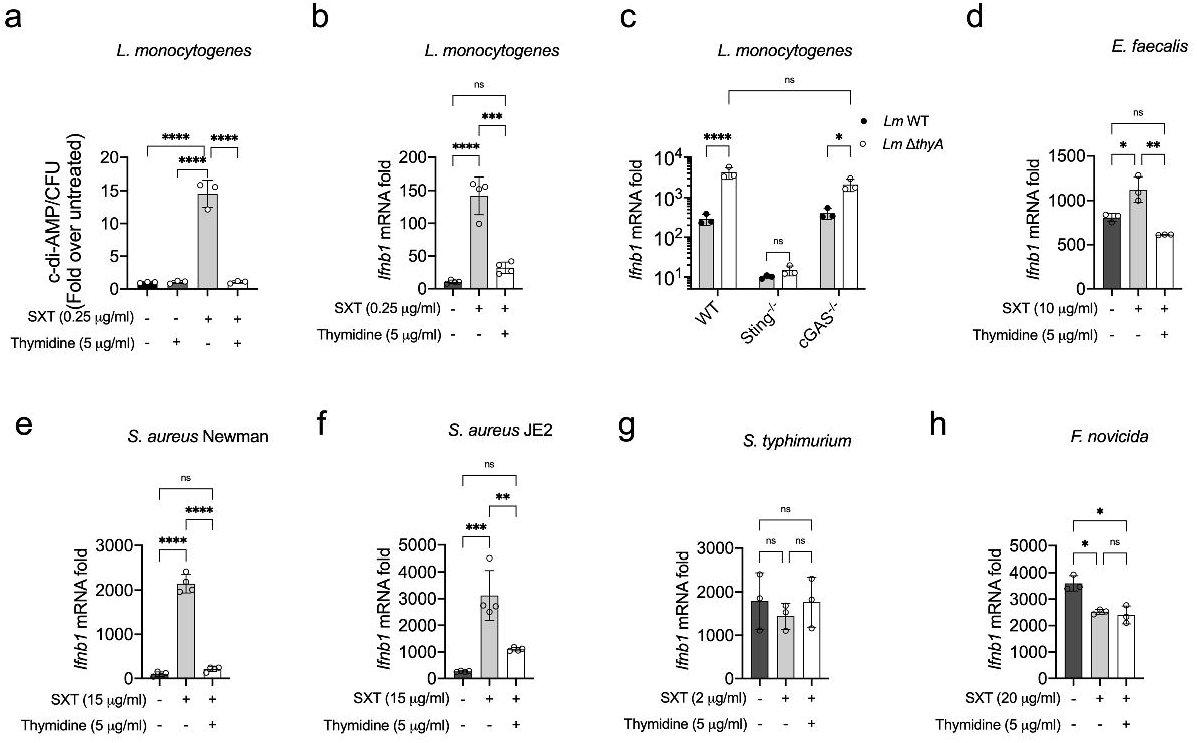
*L. monocytogenes* ThyA inhibition or mutation leads to higher c-di-AMP production. **a**, Intracellular c-di-AMP concentration of *Listeria monocytogenes* grown in BHI broth with indicated SXT and thymidine supplementation. **b**, *Ifnb1* mRNA induction in pBMDMs infected with *L. monocytogenes* grown in BHI broth as indicated in (**a**). **c**, *Ifnb1* mRNA induction by WT or Δ*thyA L. monocytogenes* strains in WT, *Sting* ^-/-^ or *cGas*^-/-^ pBMDMs at 4 hpi. **d**-**h**, *Ifnb1* mRNA induction in pBMDMs infected with either *Enterococcus faecalis* (**d**), *S. aureus* Newman (**e**), *S. aureus* JE2 (**f**), *Salmonella enterica* serovar Typhimurium (**g**) and *Francisella novicida* (**h**) grown in the indicated conditions. For all panels, mean values of biological replicates are plotted, and error bars indicate ±SD, *P* values were calculated using Two-way ANOVA analysis. Asterisks indicate that differences are statistically significant (*, *P* < 0.05, **, *P* < 0.01; ****, *P* < 0.0001), and ns indicates no significant difference.

### *S. aureus* TD-SCVs activate STING signaling through c-di-AMP

Our findings reveal a conserved connection between disruption of thymidine synthesis and STING-dependent pro-inflammatory activity among Firmicutes. Folate synthesis is essential for the growth of bacteria, and SXT is widely used as a common medication to treat various bacterial infections, including the pulmonary, skin, and soft tissue infections caused by *S. aureus*. However, the use of SXT can result in the emergence of antifolate-resistant *S. aureus*^32,33^, referred to as thymidine-dependent small-colony variants (TD-SCVs). TD-SCVs are characterized by thymidine auxotrophy due to inactivating mutations in *thyA*, yielding non-hemolytic, small colonies when cultured on blood agar plates. Clinically, TD-SCVs are associated with chronic and recurring antibiotic resistant infections. Notably, chronic respiratory infection with TD-SCVs was recently found to be associated with significantly increased risk of respiratory exacerbations and reduced lung function in pediatric CF patients, whereas those with non-thymidine-dependent SCVs did not^27,28^. While the link between TD-SCVs and worse lung disease has been well documented^27,28,34^, the mechanism behind these observations is not understood. We hypothesized that this clinical association resulted from increased inflammation induced by TD-SCV infections relative to other *S. aureus* isolates. In support of this hypothesis, *Ifnb1* expression following macrophage infection was similarly induced by methicillin-susceptible *S. aureus* (MSSA, strain Newman), and methicillin-resistant *S. aureus* (MRSA, strain JE2) after SXT treatment in the presence or absence of thymidine supplementation conditions (Fig. 2e, f) that phenocopy the effects of TD-SCV mutations. We therefore further characterized the impacts of SXT and Δ*thyA* mutation on the dynamics of infection by *S. aureus* and resulting inflammation, focusing on clinical isolates of *S. aureus*.

We obtained two sets of clonally-related *S. aureus* normal colony (NC) and SCV isolates collected during a study of children with CF^35^ and infected pBMDMs with each. Consistent with our prediction, we found that the clinical TD-SCV isolates exhibited between 10 and 100-fold higher levels of c-di-AMP/CFU (Fig. 3a) and simultaneously induced significantly higher *Ifnb1* in both WT and *cGas*^-/-^ pBMDMs relative to normal-colony (NC) clonal isolates (Fig. 3b). These observations suggest that the macrophage inflammatory response is elevated during *S. aureus* infection under two conditions that limit thymidine levels: following SXT treatment, and when *S. aureus* carries inactivating mutations in the *thyA* gene.

**Fig. 3:**
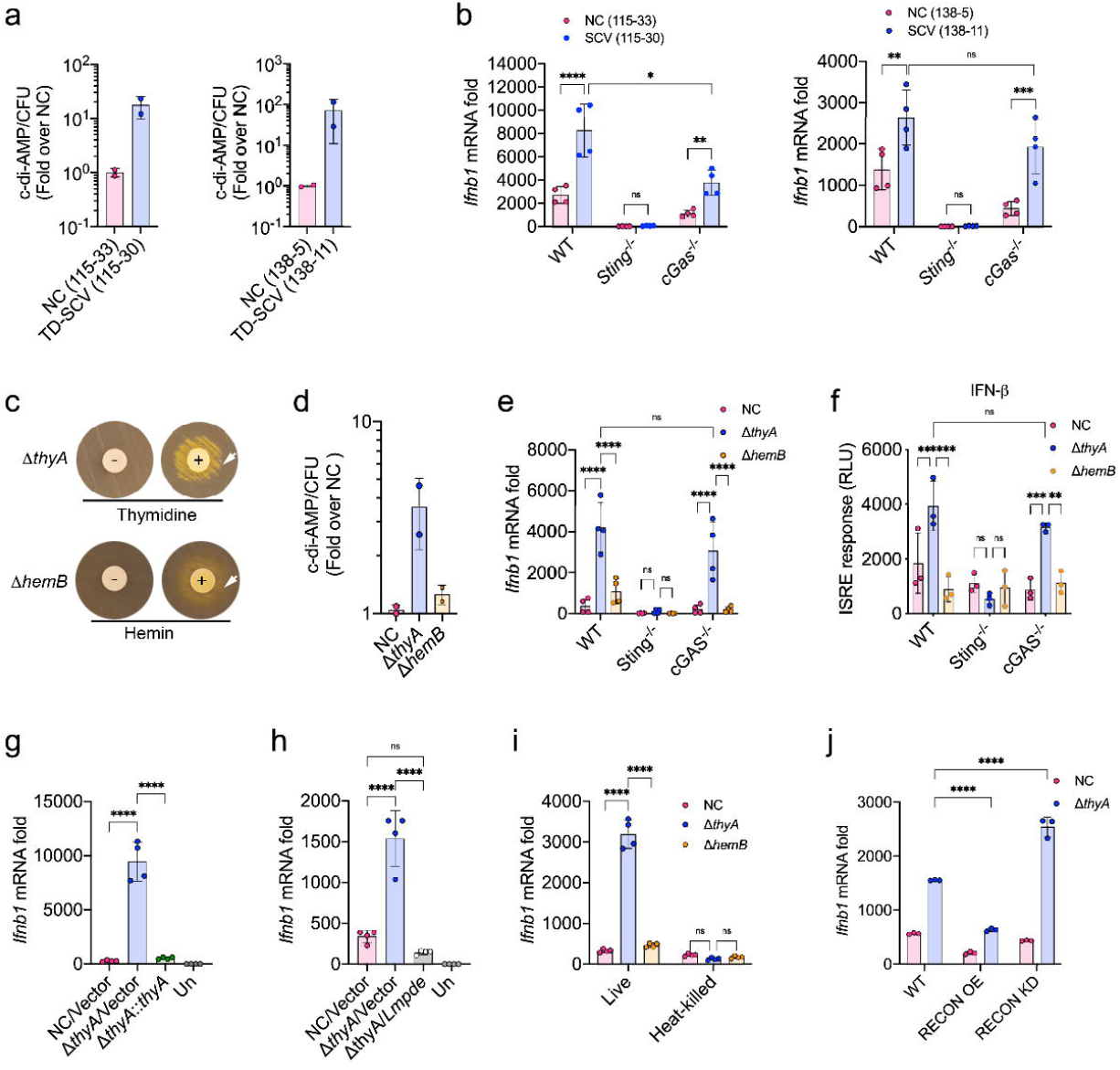
Δ*thyA* activates STING-dependent type I IFN production through c-di-AMP in macrophages. **a**, LC-MS/MS detection of c-di-AMP levels normalized for cell density after overnight growth on chocolate agar by clinical TD-SCV isolates (115-30 and 138-11) relative to their clonally-related normal-colony (NC) counterpart isolates (115-33 and 138-5). **b**, *Ifnb1* expression in iBMDMs infected with clinical TD-SCV isolates at 4 hpi relative to uninfected control. **c**, Auxotrophy test for thymidine and hemin dependence in the deletion strains. Representative image of a lawn of Δ*thyA* or Δ*hemB* plated on Mueller-Hinton agar and showing enhanced growth around paper disks impregnated with thymidine or hemin, respectively. The arrows indicate bacterial growth around the disk. **d**, LC-MS/MS quantification of c-di-AMP levels of Δ*thyA* and Δ*hemB* relative to NC. **e**, WT, *Sting* ^-/-^ and *cGas*^-/-^ pBMDMs were infected with NC, Δ*thyA* and Δ*hemB* Newman strains for 4 hrs, and induction of *Ifnb1* transcripts were measured by qRT-PCR relative to uninfected controls (Un) incubated for the same duration. **f**, IFN-β protein concentration in pBMDM supernatants was measured by ISRE-luciferase bioassay at 6 hpi. **g**, WT iBMDMs were infected with NC, Δ*thyA* and Δ*thyA::thyA* Newman strains for 4 hrs, and *Ifnb1* mRNA was measured by qRT-PCR relative to uninfected controls. **h**, *Ifnb1* expression in iBMDMs infected with NC, Δ*thyA*, and *L. monocytogenes pde* (*Lmpde*) overexpression Newman strains relative to uninfected control. Mean values of 4 replicates are plotted and error bars indicate ±SD. **i**, *Ifnb1* transcription induction in iBMDMs by live or heat-killed NC, Δ*thyA* and Δ*hemB* Newman strains with the same starting CFU assayed at 4 hpi. **j**, *Ifnb1* expression in WT, RECON overexpression (RECON OE) or knockdown (RECON KD) iBMDMs infected with NC or Δ*thyA* Newman strains relative to uninfected controls. All infections were performed at an MOI of 20. For all panels, mean values of biological replicates are plotted, and error bars indicate ±SD, *P* values were calculated using Two-way ANOVA analysis. Asterisks indicate that differences are statistically significant (*, *P* < 0.05; **, *P* < 0.01; ***, *P* < 0.001, ****, *P* < 0.0001), and ns indicates no significant difference.

While our comparison of the effects of clinical TD-SCV isolates with those of clonally-related normal-colony isolates indicates *thyA* mutation was responsible for our observations, it was possible that the clinical TD-SCVs also carried secondary mutations that influence the c-di-AMP production or host IFN induction during infection. We therefore generated a laboratory TD-SCV strain by deleting the *thyA* gene in *S. aureus* Newman. Additionally, we constructed a hemin-dependent SCV (HD-SCV) control strain containing a deletion in the porphobilinogen synthase gene *hemB* in *S. aureus* Newman. Δ*hemB* was selected as an SCV control to compare to the Δ*thyA* strain because HD-SCVs are also observed during human infections, and although hemin-deficient SCVs exhibit slow growth rates and antibiotic resistance, they are not known to be associated with worse CF lung disease outcomes^27^.

We confirmed that the growth of Δ*thyA* and Δ*hemB* on indicator media was complemented by supplementation of exogenous thymidine or hemin, respectively (Fig. 3c). At logarithmic phase, Δ*thyA* produced about 2-fold higher c-di-AMP than both NC and Δ*hemB* strains grown in BHI broth, even with surplus thymidine supplementation (5 µg/ml) (Fig. 3d), which phenocopied the clinical TD-SCVs (Fig. 3a). Next, WT, *Sting*^-/-^ and *cGas*^-/-^ pBMDMs were infected with either NC, Δ*thyA* or Δ*hemB S. aureus* Newman strains, and intracellular bacterial survival was measured under each condition. NC and Δ*hemB* exhibited similar survival trends over 8 hours post infection (hpi), whereas Δ*thyA* exhibited significantly reduced persistence in all cell lines (Extended Data Fig. 1a). Additionally, all strains generated comparably low levels of macrophage toxicity, as assessed by LDH release (Extended Data Fig. 1b). Despite the diminished CFU burden, *S. aureus* Δ*thyA* induced strikingly higher STING-dependent and cGAS-independent *Ifnb1* transcription and IFN-β levels than either NC or Δ*hemB* strains (Fig. 3e-f).

Complementation of Δ*thyA in trans* with the *thyA* gene driven by its original promoter exhibited similar *Ifnb1* induction as the WT Newman strain (Fig. 3g) and rescued the survival defect of Δ*thyA* within cells (Extended Data Fig. 1c). Unlike the clinical TD-SCV isolate 115-30, which elicited noticeable cGAS-dependent *Ifnb1* expression (Fig. 3b), *Ifnb1* induction by Δ*thyA* was totally abolished in *Sting*^-/-^ pBMDMs but only slightly diminished in *cGas*^-/-^ pBMDMs (Fig. 3e), consistent with the observations for the Δ*thyA L. monocytogenes* strain (Fig. 2c).

Bacteria are known to produce a diverse range of cyclic dinucleotides, including c-di-GMP, c-di-AMP, and 3′3′-cGAMP, among others^36,37^. However, *S. aureus* Newman is reported to lack functional enzymes to synthesize either 3′3′-cGAMP^36^ or c-di-GMP^38,39^, and this strain also lacks alternative CD-NTase enzymes to synthesize other cyclic dinucleotides^37^, suggesting that c-di-AMP is the primary cyclic dinucleotide capable of activating STING produced by this strain. To further verify that the STING activation upon Δ*thyA* infection is due to c-di-AMP, Δ*thyA* overexpressing the *L. monocytogenes* phosphodiesterase PdeA, which degrades c-di-AMP^40^, was used for macrophage infection along with NC and Δ*thyA* strains. Consistent with our hypothesis, expression of the c-di-AMP phosphodiesterase in Δ*thyA* eliminated its *Ifnb1* induction (Fig. 3h). Furthermore, c-di-AMP represents a key viability-associated pathogen-associated molecular pattern (vita-PAMP) capable of activating STING to orchestrate host immune responses to live bacteria^41^. Heat-killing of bacterial cells abolished c-di-AMP production and consequently blunted *Ifnb1* induction (Fig. 3i). We also investigated the effects of c-di-AMP sequestration on *Ifnb1* induction by manipulating the expression of the oxidoreductase RECON, which was identified previously by our laboratory as a high-affinity host receptor specific for bacterial cyclic dinucleotides^42^. RECON antagonizes STING activation by acting as a molecular sink for c-di-AMP during bacterial infection in macrophages. To determine the effect of RECON overexpression on *Ifnb1* induction during Δ*thyA* infection, the WT, RECON overexpression (RECON OE) and knockdown (RECON KD) immortalized bone marrow derived macrophages (iBMDMs) were each infected with NC and Δ*thyA S. aureus* Newman (Fig. 3j). As expected, RECON overexpression disrupted *Ifnb1* induction by Δ*thyA*, whereas RECON knockdown significantly increased *Ifnb1* expression. These results further demonstrate that excessive c-di-AMP production by *S. aureus* Δ*thyA* is responsible for STING activation during infection.

### Thymidine availability influences c-di-AMP production by TD-SCVs

Mutations in *thyA* disrupt dTMP (deoxythymidylate) synthesis from dUMP (deoxyuridylate), resulting in reliance on exogenous thymidine to maintain DNA synthesis^43^. We observed that exogenous thymidine supplementation dampened the IFN-β induction by c-di-AMP producing Firmicutes *L. monocytogenes* (Fig. 2b), *E. faecalis* (Fig. 2d), and *S. aureus* strains Newman (Fig. 2e) and JE2 (Fig. 2f) when treated with SXT. As such, we sought to investigate the effect of *thyA* deletion and exogenous thymidine availability on bacterial c-di-AMP production. Supplementation of thymidine restored the growth defect of Δ*thyA* in BHI broth in a dose-dependent fashion (Fig. 4a). We then determined the effect of thymidine supplementation on c-di-AMP production. Relative to NC *S. aureus*, intracellular c-di-AMP levels of Δ*thyA* were markedly increased upon thymidine starvation (1.25 µg/mL) compared with thymidine concentrations (2.5 and 5 µg/mL) that maximized growth (Fig. 4b), indicating that thymidine availability regulates c-di-AMP production when *thyA* is inactivated. We quantified the c-di-AMP levels in macrophages infected with logarithmic-phase NC, Δ*thyA* or Δ*hemB S. aureus* strains. Consistent with our previous data, macrophages infected with Δ*thyA* had higher c-di-AMP levels relative to WT or Δ*hemB* infected cells (Fig. 4c). We also determined c-di-AMP levels in macrophages infected with Δ*thyA* or the NC strain grown in a range of thymidine concentrations, and measured *Ifnb1* mRNA levels in parallel (Fig. 4d). Macrophages infected with Δ*thyA* grown in lower concentrations of thymidine exhibited higher c-di-AMP levels and a correspondingly elevated *Ifnb1* transcriptional response, whereas both c-di-AMP and *Ifnb1* mRNA levels in macrophages infected with WT did not vary when grown in different thymidine concentrations (Fig. 4e-f). These observations suggest that TD-SCV infection leads to higher STING activation than NC *S. aureus*, which is exacerbated upon thymidine limitation.

**Fig. 4:**
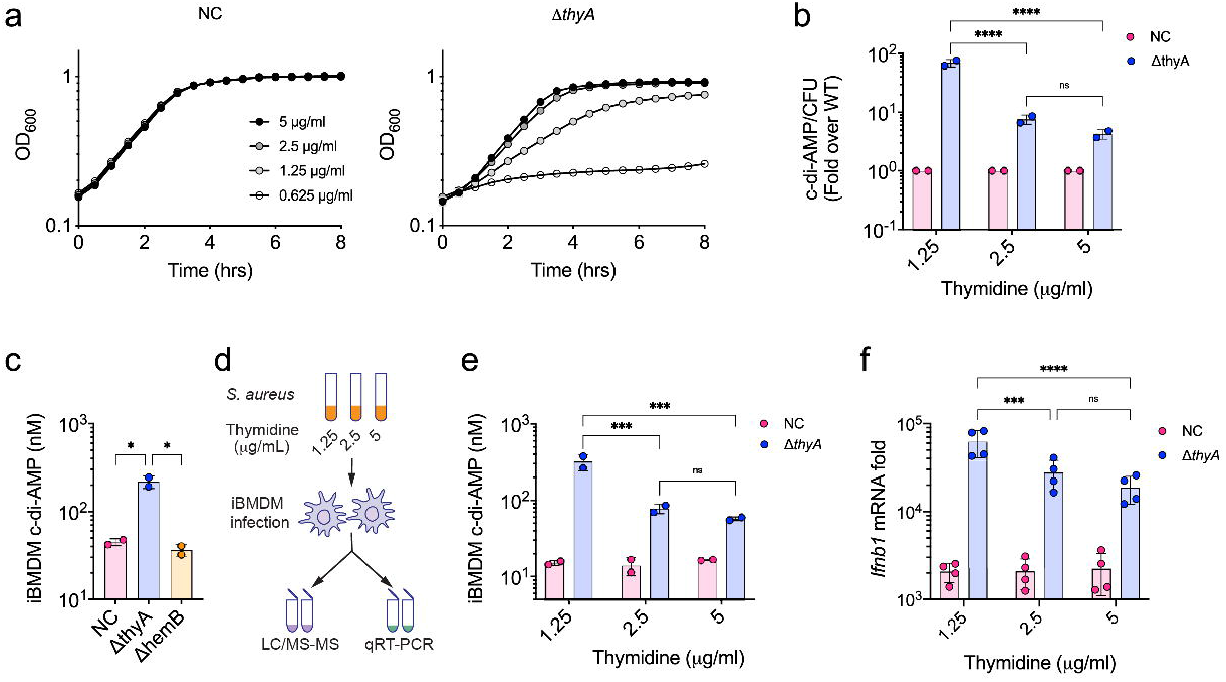
Thymidine abundance modulates c-di-AMP levels and STING-activating capacity of TD-SCV. **a**, Growth curve of NC (left panel) and Δ*thyA* (right panel) *S. aureus* Newman in BHI broth supplemented with the indicated concentrations of thymidine. **b**, LC-MS/MS quantitation of c-di-AMP levels normalized for CFU, of NC and Δ*thyA* grown in BHI broth supplemented with the indicated concentrations of thymidine. **c**, LC-MS/MS quantification of intracellular c-di-AMP in iBMDMs infected with NC, Δ*thyA* and Δ*hemB* Newman strains for 4 hrs. **d**, Schematic diagram of macrophage infection, in which WT iBMDMs were infected with NC or Δ*thyA* overnight culture grown in BHI broth supplemented with the indicated thymidine concentrations for 4 hrs, the intracellular c-di-AMP levels and *Ifnb1* expression in iBMDMs were determined by LC-MS/MS and qRT-PCR, respectively. **e**, The intracellular c-di-AMP concentration in iBMDMs as in panel **d**). **f**, Infections performed as in **d**) and *Ifnb1* mRNA was measured by qRT-PCR relative to uninfected control. All the infections were performed at the MOI of 20. For all panels, mean values of biological replicates are plotted, and error bars indicate ±SD. *P* values were calculated using One or Two-way ANOVA analysis. Asterisks indicate that differences are statistically significant (**, *P* < 0.01; ****, *P* < 0.0001).

### Airway TD-SCV infection elicits neutrophil infiltration through STING activation

The association with TD-SCVs and declining lung function among CF patients is unexpected given their diminished capacity to survive within host phagocytes and previous reports demonstrating their diminished virulence^44^ (Extended Data Fig. 1). Given our findings that elevated c-di-AMP production by TD-SCVs promotes increased STING-dependent inflammation, we were intrigued to explore the impacts of this pathway in an *in vivo* context. Taqman cytokine arrays in macrophages infected with TD-SCV revealed significant elevation in the production of the IFN-β co-regulated cytokines IL-6, CXCL10 and CCL5 (Extended Data Fig. 2), which are expressed upon activation of STING signaling^10,45,46^. To interrogate the role of c-di-AMP in promoting airway inflammation, we quantified levels of these cytokines in the bronchoalveolar lavage fluid (BALF) collected from WT or *Sting*^-/-^ mice following intranasal instillation with the TLR2 agonist Pam3CSK4 either alone or in combination with c-di-AMP. Induction of these cytokines by Pam3CSK4 and c-di-AMP was found to be both STING and c-di-AMP dependent (Fig. 5a-c). Notably, these cytokines are known for mediating recruitment and activation of neutrophils, the primary innate immune cell type in the defense against *S. aureus* infections^47^, as well as neutrophil-mediated pulmonary inflammation and injury^48–50^. Cell counts of the BALF revealed significant c-di-AMP-dependent elevation of neutrophil migration into the airway (Fig. 5d).

**Fig. 5:**
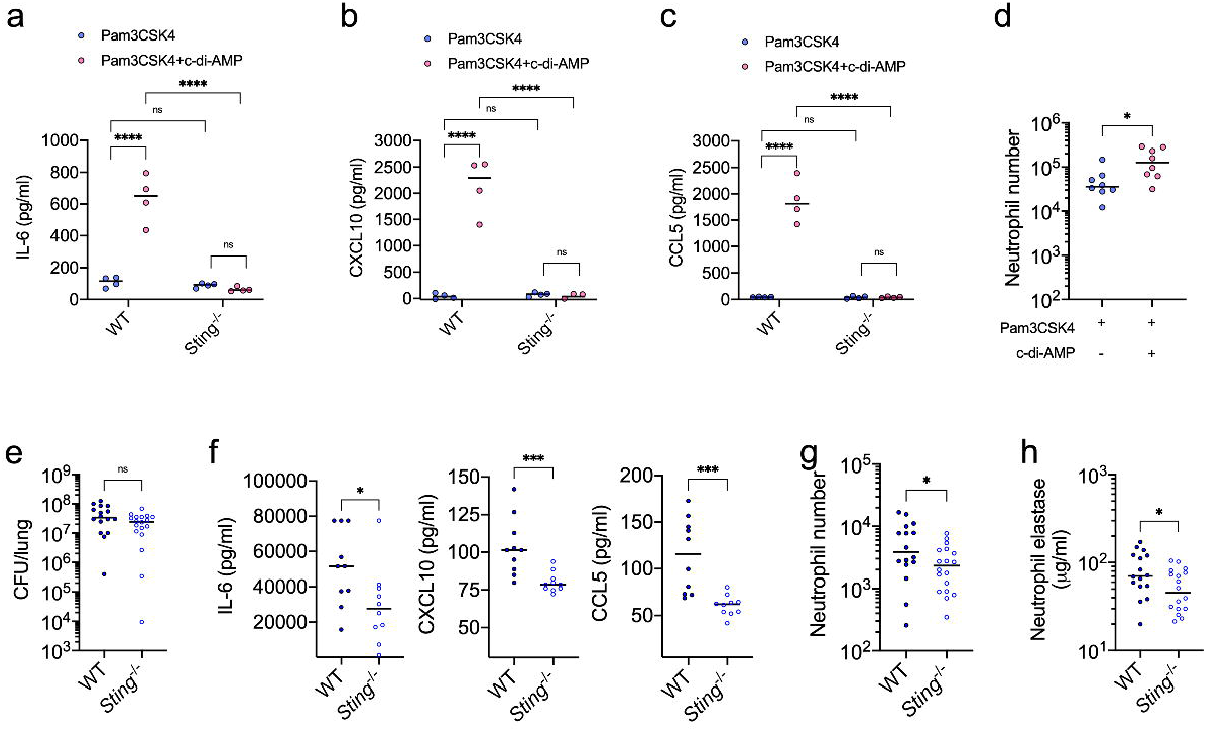
STING controls the airway neutrophil activation induced by TD-SCV infection. **a**-**c**, The concentration of cytokine IL-6 (**a**), CXCL10 (**b**), and CCL5 (**c**) in BALF collected from the WT or *Sting*^-/-^ mice which intranasally instilled with 15 μg Pam3CSK4 and 15 μg c-di-AMP or Pam3CSK4 alone as quantified by ELISA. **d**, The neutrophils present in BALF collected from the WT mice intranasal instillation with 15 μg Pam3CSK4 and 15 μg c-di-AMP or Pam3CSK4 alone as identified and enumerated by FACS assay. **e**, The CFU recovery of Δ*thyA* from the right lungs of WT or *Sting*^-/-^ mice at 8 hpi. WT or *Sting*^-/-^ mice were intranasally infected with Δ*thyA* strain with 5×10^8^ CFU/mouse. The BALF was collected and the CFU recovery from the right lung were enumerated at 8 hpi. **f**, BALF IL-6, CXCL10 and CCL5 concentration of mice infected as panel (**e**) determined by ELISA. **g**, The neutrophils present in BALF of WT or *Sting*^-/-^ mice with Δ*thyA* infection as identified and enumerated by FACS assay. **h**, Active neutrophil elastase concentration in BALF from mice infected with Δ*thyA*. For all the mouse infections, biological replicates are plotted, and horizontal black bars are the median of the data.*P* values were calculated using Mann-Whitney analysis. Asterisks indicate that differences are statistically significant (*, *P* < 0.05; **, *P* < 0.01; ***, *P* < 0.001), and ns indicates no significant difference.

Given that TD-SCVs produce excessive c-di-AMP, we hypothesized that TD-SCV infection could induce airway neutrophil infiltration similar to the effects of c-di-AMP. To investigate this possibility, WT and *Sting*^-/-^ mice were intranasally infected with the Δ*thyA* strain followed by bronchoalveolar lavage and lung CFU enumeration at 8 hpi. Consistent with the tissue culture studies (Extended Data Fig. 1a), *Sting* deletion did not affect bacterial survival at 8 hpi (Fig. 5e), however, the production of IL-6, CXCL10 and CCL5 was significantly decreased in *Sting*^-/-^ mice (Fig. 5f), which is consistent with the results from c-di-AMP instillation (Fig. 5a-c). Additionally, *Sting*^-/-^ mice demonstrated significantly decreased airway neutrophil infiltration (Fig. 5g). The active neutrophil elastase concentration in BALF, which reflects activated neutrophil levels, was also significantly higher in WT than *Sting*^-/-^ mice (Fig. 5h). These observations are consistent with elevated c-di-AMP production in Δ*thyA* strains resulting in STING-dependent inflammation and subsequent neutrophil migration into the airway during lung infection.

### Recurrent TD-SCV infection induces higher airway inflammation than other *S. aureus* subtypes

Having demonstrated that TD-SCVs promote increased STING-dependent inflammation and neutrophil recruitment in the lung, we subsequently determined if these observations were unique to TD-SCVs relative to NC and Δ*hemB S. aureus* strains. As expected, in WT mice, Δ*thyA* infection resulted in approximately 10-fold lower bacterial recovery from lung tissue (Fig. 6a) but induced significantly higher production of c-di-AMP-STING-dependent cytokines IL-6, CXCL10 and CCL5 in the BALF relative to NC strains at 8 hpi (Fig. 6b). Mice infected with Δ*thyA* also exhibited 10-fold elevation of airway neutrophil infiltration (Fig. 6c), and significantly higher active neutrophil elastase concentration in the BALF relative to NC infection (Fig. 6d). We also revealed that Δ*thyA* was recovered with diminished abundance (Extended Data Fig. 3a, Extended Data Fig. 4) from lung tissue but induced more neutrophil infiltration (Extended Data Fig. 3b), higher neutrophil elastase activity (Extended Data Fig. 3c) and STING-dependent cytokines (Extended Data Fig. 3d-f) compared with the Δ*hemB* strain. These observations establish that TD-SCVs are characterized by elevated inflammatory capacity relative to normal colony *S. aureus* or SCVs of other genotypes.

**Fig. 6:**
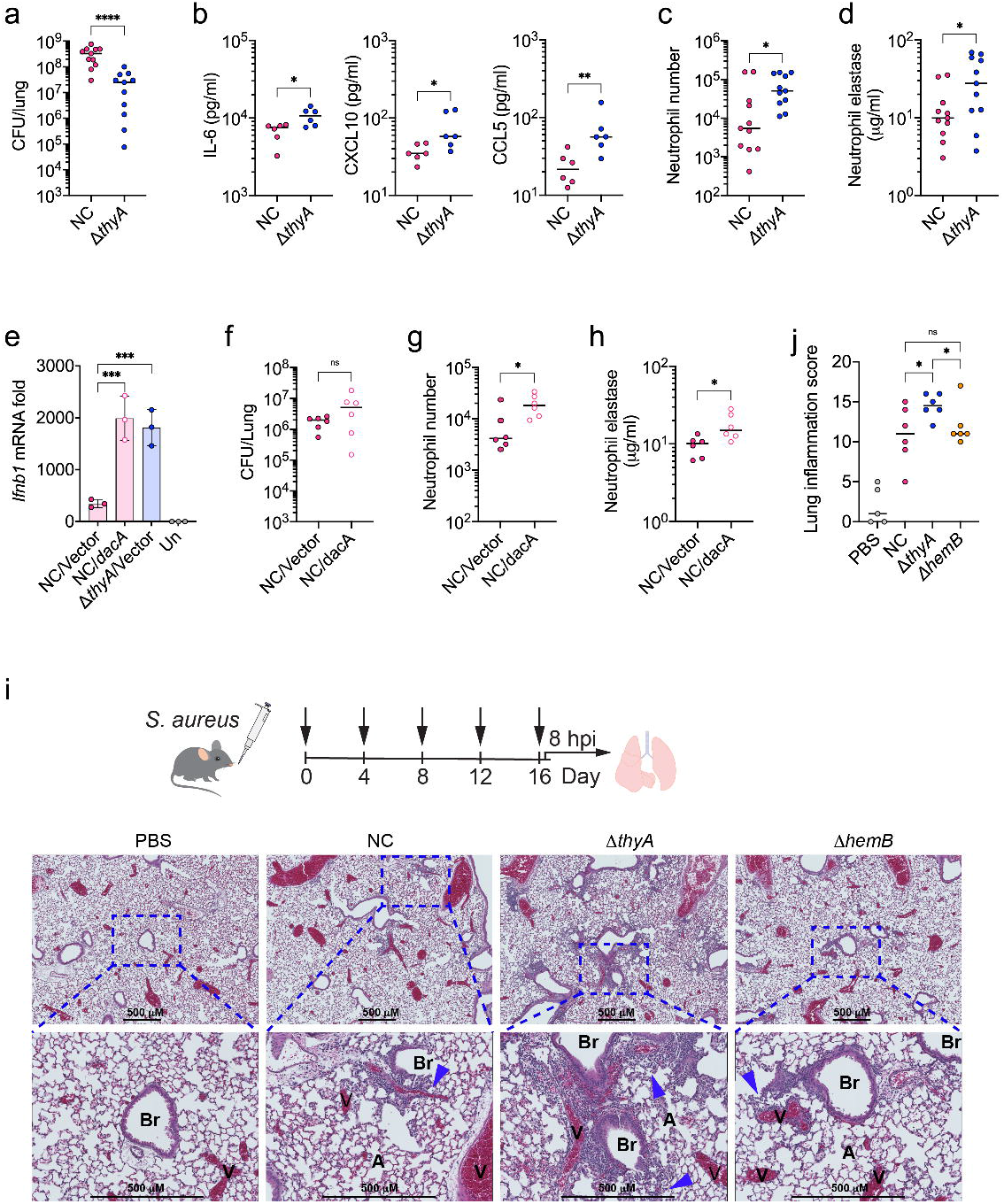
TD-SCV induces robust airway neutrophil infiltration and inflammation during both acute and chronic infection. **a**, CFU recovery of either NC or Δ*thyA* strain from WT mice. **b**, BALF IL-6, CXCL10 and CCL5 concentration of mice infected as panel (**a**) determined by ELISA.**c**, The neutrophils in BALF of the mice infected as in panel (**a**) were identified and enumerated by FACS assay. **d**, Active neutrophil elastase concentration in BALF from mice infected as in panel (**a**). **e**, *Ifnb1* expression in iBMDMs infected with NC, *dacA* overexpression, or Δ*thyA* strains relative to uninfected control. Mean values of triplicates are plotted and error bars indicate ±SD. *P* values were calculated using Two-way ANOVA analysis. **f**, WT mice were intranasally infected with *dacA* overexpression (NC/*dacA*) or its control (NC/vector) Newman strain. The CFU recovery from the right lung were enumerated at 8 hpi. **g**, The neutrophils present in BALF collected from the mice infected as in panel (**f**) were identified and enumerated by FACS assay. **h**, Active neutrophil elastase concentration in BALF from the mice infected as in panel (**f**). **i**, The top panel indicates the repeat infection strategy, in which WT mice were intranasally instilled with either NC, Δ*thyA* or Δ*hemB* strain with 2.5×10^7^ CFU/mouse or PBS every 4 days for a total of 5 infections. The lungs were harvested 8 hrs after the last infection. The bottom panel indicates the representative images of H&E staining of WT mouse lung sections. The arrows indicate the inflammatory cell infiltrate. Br = bronchiole, V = blood vessel, and A = alveoli. **j**, Inflammation scoring for mice infected as (**i**). The inflammation scores were the summation of perivascular/peribronchiolar, intrabronchiolar, intraalveolar, interstitial inflammation and overall severity. For all mouse infection panels, biological replicates are plotted. Horizontal black bars indicate the median of the data. *P* values were calculated using Mann-Whitney analysis. Asterisks indicate that differences are statistically significant (*, *P* < 0.05; **, *P* < 0.01; ****, *P* < 0.0001), and ns indicates no significant difference.

To further confirm if excessive c-di-AMP production in *S. aureus* led to the higher inflammation and neutrophil recruitment, a *S. aureus* Newman strain ectopically overexpressing the di-adenylate cyclase DacA (NC/*dacA*), which synthesizes c-di-AMP^51^, was generated. STING activation in macrophages was confirmed along with the control strain (NC/vector) by measuring *Ifnb1* expression (Fig. 6e). NC *S. aureus* p::*dacA* exhibited similar bacterial survival relative to control bacteria (Fig. 6f) but induced significantly higher neutrophil infiltration and activation as assessed by active elastase concentration in WT mice at 8 hpi (Fig. 6g-h). This finding establishes that elevated c-di-AMP production is sufficient to promote neutrophil recruitment into the airway during infection.

The lungs of CF patients are characterized by thick mucus build up, leading to an environment that favors chronic bacterial infection. Chronic and recurrent *S. aureus* infection causes lung inflammation and permanent lung damage of patients with CF despite antibiotic intervention. Based on our observations, we hypothesized that the hyperinflammatory capacity of TD-SCVs could promote respiratory inflammation in chronic or recurrent infections. To determine whether the higher inflammation that occurred during Δ*thyA* infection also resulted in associated pathology during long term infection, we performed semi-quantitative histopathological analysis of the lungs from mice intranasally instilled with either NC, Δ*thyA* or Δ*hemB* strains or PBS in a repeat infection model. After five infections total over sixteen days, mice infected with all tested *S. aureus* strains exhibited robust airway inflammatory cell infiltration compared to mice treated with PBS (Fig. 6i), whereas mice with Δ*thyA* infection exhibited elevated inflammatory cell infiltration and significantly higher inflammation scores relative to the NC and Δ*hemB* strains (Fig. 6j, Supplemental Table 1). These observations establish that despite their diminished capacity to survive *in vivo*, TD-SCVs exhibit elevated inflammatory capacity that promotes immune cell migration to the airway during acute infection. These observations may similarly explain the observed association of TD-SCVs with worse respiratory outcomes during chronic infection.

## Discussion

The results of this study clearly demonstrate that disruption of thymidine metabolism results in the elevated production of the second messenger c-di-AMP within several bacterial species, implicating this signaling molecule as a mediator of the stress imposed by thymidine limitation. In addition to its role as a key signaling molecule within bacteria, c-di-AMP also functions as a potent activator of STING-dependent inflammation within mammalian hosts^2^. Indeed, our findings demonstrate that inhibition of thymidine metabolism by commonly-used antibiotics significantly increased STING-dependent inflammatory responses following bacterial infection. Additionally, antifolate-resistant thymidine auxotrophs of *S. aureus*, which are frequently isolated from CF patients and are associated with lower lung function among this patient population, similarly exhibit increased STING-dependent inflammation and increased inflammatory cell recruitment into the airway relative to other (less common) SCV types or normal colony *S. aureus*.

To date, no link between bacterial c-di-AMP production and thymidine starvation has been reported. While the molecular mechanism behind this observation remains unresolved, it suggests that c-di-AMP signaling may be involved directly in countering the stress associated with thymidine limitation. For instance, thymidine depletion is known to induce DNA damage^52^, and c-di-AMP signaling has been implicated in the repair of DNA lesions^53^. It is therefore tempting to speculate that c-di-AMP levels in TD-SCVs are modulated by pyrimidine metabolism, perhaps through an effect of thymidine starvation on DNA repair pathways. Alternatively, c-di-AMP is a general osmotic stress regulator involved in bacterial uptake of potassium and compatible solutes that function to maintain cellular turgor. Modulation of c-di-AMP by *S. aureus* impacts cell size and moderates the effects of extreme membrane and cell wall stress by altering cell size and the composition of the cell wall^38,54^. Cells of Δ*thyA S. aureus* are abnormally shaped during thymidine limitation^44^ and c-di-AMP levels may be modulated in response to distorted cellular structure, perhaps ensuring cell integrity.

While it is conceivable that c-di-AMP impacts cellular physiology following thymidine limitation, other observations suggest a different relationship may link these processes. The DNA synthesis precursor dTMP can be generated either by reductive methylation of dUMP by ThyA, or by phosphorylation of thymidine by thymidine kinase (TMK)^43^. Consequently, disruption of *thyA* causes bacterial reliance on exogenous thymidine for growth. In *L. monocytogenes, E. faecalis*, and *S. aureus*, the gene *tmk* is in a conserved operon with the c-di-AMP receptor, PII-like signal transduction protein A (*pstA*). PstA has no biological function described and it is intriguing to speculate that it may affect TMK activity^55,56^. Future studies to delineate the molecular mechanism linking c-di-AMP and thymidine starvation will shed significant insight into the role of this second messenger in the physiology of bacteria.

Antibiotic therapy influences the inflammatory properties of bacteria by promoting release of inflammatory components^57,58^. In many cases these proinflammatory effects of antibiotic exposure work in concert with the impacts on microbial growth to clear infection. In some instances, such as during treatment of infections with high bacterial burdens, excessive inflammation can promote toxic shock. Such processes are most well-described for cell wall-targeting antibiotics. Our observations suggest that antifolates may have similar beneficial and detrimental effects depending on the infection context. During acute infections, STING-dependent inflammation may augment the bactericidal activity of these antibiotics in favor of controlling infection. However, the capacity of TD-SCVs to hyperactivate STING and their association with worse lung outcomes suggests a potential role for STING activation and pathological tissue damage in a chronic infection context. Our study coupled STING activation with neutrophil infiltration and activation in response to TD-SCV infection in the airway. Neutrophils protect the host against bacterial infection by production of neutrophil extracellular traps (NETs), reactive oxygen and nitrogen species, and anti-microbial proteases and peptides^59^. However, persistent neutrophil recruitment and activation can lead to over-exuberant, and maladaptive, tissue damage^60^. Therefore, chronic inflammatory cell recruitment may damage the airway epithelium and drive lung function decline. Indeed, lung disease is a major complication of the autoimmune disease **S**TING-**A**ssociated **V**asculopathy with onset in **I**nfancy (SAVI)^17^, which is caused by STING gain-of-function mutations in humans. Future studies detailing the impacts of STING activation in the context of CF and other chronic infections associated with TD-SCVs may provide new therapeutic interventions to prevent pathology associated with their unique hyperinflammatory properties.

While STING impacts inflammatory cell recruitment during lung infection by TD-SCVs, it has not escaped our attention that our data implicate a role for other unique inflammatory properties of this SCV genotype. Specifically, neutrophil infiltration induced by Δ*thyA* infection is more than 5-fold higher than that with NC or Δ*hemB* strains, while deletion of STING decreased but did not abolish the higher neutrophil infiltration induced by Δ*thyA*. These observations indicate that Δ*thyA* also activates other STING-independent innate immune pathways that are important for sensing TD-SCV infection and recruiting neutrophils *in vivo*. Many pattern PRRs such as Toll-like (TLRs) and (NOD)-like receptors have been implicated in innate immune sensing of *S. aureus*. TLR1, 2 and 6 specifically recognize cell wall lipopeptides and peptidoglycan, whereas the intracellular PRRs, NOD1 and NOD2, detect bacterial peptidoglycan to induce inflammation^61,62^. TD-SCVs are reported to have altered cell wall composition and abnormal cell size^44,63^, which may lead to higher TLR or NOD1/2 activation, eliciting the production of inflammatory cytokines, and contributing to the hyperinflammatory capacity of TD-SCVs together with STING activation. While further work is needed to detail the mechanisms by which TD-SCV impact inflammation during infection, our studies certainly underscore the importance of considering the unique pathologies that may arise during infection with distinct SCV genotypes.

In summary, our study revealed a previously unrecognized impact of thymidine limitation on the production and immune-activating potential of Firmicutes that produce c-di-AMP. We anticipate that the findings of this study will provide insight into the physiological function of the widespread bacterial second messenger c-di-AMP and provide a broader understanding of the impacts of antifolate treatment on the immune response in the context of acute and chronic bacterial infection with bacteria that produce c-di-AMP.

## Methods

### Microbe Strains and Culturing Conditions

Clinical *S. aureus* isolates were grown on chocolate II agar (BD, USA). *L. monocytogenes* 10403s, *S. aureus* Newman and JE2 and *E. faecalis* OG1RF were grown in brain heart infusion (BHI) broth and incubated at 37 °C with shaking. *Escherichia coli* strains used for cloning and *S*. Typhimurium SL1344 were grown in Lysogeny broth (LB) at 37 °C with shaking. *F. novicida* was grown in TSBC (30 g tryptic soy broth, 1 g cysteine per liter) at 37 °C with shaking. Hemin-dependent and thymidine-dependent SCVs were grown in BHI broth supplemented with hemin or thymidine at the indicated concentration.

### Murine Cell Lines

Bone marrow from WT, *cGas*^-/-^, and *Tmem173*^-/-^ mice were generously provided by Daniel Stetson at the University of Washington. Primary BMDMs were generated as previously described ^64^. J2 virus immortalized BMDM cell line was from Dan Portnoy at the University of California, Berkeley. Both the primary and immortalized BMDMs were cultured in Dulbecco’s Modified Eagle Medium (DMEM) supplemented with 2 mM sodium glutamine, 1 mM sodium pyruvate, 10% heat-inactivated FBS, 0.1% 2-Mercaptoethanal, 10% L929 cell supernatants as a source of M-CSF and 1% penicillin/streptomycin.

### Mice

All the mice used in this study were on C57BL/6J background. The original breeding pairs of WT were from Jackson Laboratories, *Tmem173*^-/-^ were donated by Daniel Stetson at the University of Washington ^65^. Mice colonies were bred and maintained under SPF conditions and ensured through the rodent health monitoring program overseen by the Department of Comparative Medicine at the University of Washington. All experiments involving mice were performed in compliance with guidelines set by the American Association for Laboratory Animal Science (AALAS) and were approved by the Institutional Animal Care and Use Committee (IACUC) at the University of Washington. All experiments were carried out with mice aged 8-12 weeks matched by gender, age, and body weight.

### Evaluation of inflammatory capacity of bacteria after antibiotic treatment by BIOLOG^™^ PM microplates

50 μL of *L. monocytogenes* suspension in BHI broth with an initial OD_600_ of 0.25 was added to BIOLOG™ microplates (PM11C and PM12B) or a clean 96-well cell suspension plate as an untreated control. The plates were sealed using Breathe-Easy film (Diversified Biotech, USA) to allow exchanging of oxygen and statically incubated in 37 °C for 6 hrs. The pBMDMs (5×10^5^ cells/well) were plated in 96-well tissue culture plates and infected with *L. monocytogenes* at a multiplicity of infection (MOI) of 4. *L. monocytogenes* was washed off by PBS at 1 hpi, and the cells were either lysed for CFU enumeration or continue incubating in DMEM medium with 50 µg/mL gentamicin. At 6 hpi, 25 μL cell suspensions were added to IFN responsive ISRE-L929 cells (5×10^4^ cells/well) in 96-well TC plates and incubated for 12 hrs. The media was then aspirated, and 50 μL of TNT lysis buffer (20 mM Tris, 100 mM NaCl, 1% Triton X-100) was added to each well. 40 μL of the lysate was mixed with 40 μL of luciferase substrate solution (20 mM Tricine, 2.67 mM MgSO_4_7H_2_O, 0.1 mM EDTA, 33.3 mM DTT, 530 μM ATP, 270 μM acetyl CoA lithium salt, 470 μM luciferin, 5 mM NaOH, 265 μM magnesium carbonate hydroxide) and luminescence was measured using a Synergy HT (BioTek, USA). For confirmation of the phenotype from BIOLOG™ microplates, penicillin G, 2,4-Diamino-6,7-diisopropylpteridine, sulfamethoxazole, trimethoprim, SXT (trimethoprim and sulfamethoxazole combination in a 1:5 ratio), and novobiocin was applied in various dilutions to *L. monocytogenes* suspension.

### Macrophage infection with bacteria

For macrophage infection of bacteria treated by SXT, overnight culture of *L. monocytogenes* 10403s, *E. faecalis* OG1RF, *S. aureus* Newman, *S. aureus* JE2 in BHI broth, *F. novicida* in TSBC (30 g tryptic soy broth, 1 g cystein per liter) and *S. typhimurium* (strain SL1344) in LB broth were back diluted to OD_600_ of 0.05 and incubated in various concentration of SXT overnight. 1×10^6^ of pBMDMs were plated in 6-well tissue culture plates and incubated at 37 °C in 5% carbon dioxide (CO_2_) overnight. Bacteria were washed with sterile phosphate-buffered saline (PBS) and then re-suspended in DMEM medium. The macrophages were infected by *L. monocytogenes* at a MOI of 4, *S. aureus* at a MOI of 5, *S. typhimurium* at a MOI of 1. After a 1 hr incubation, the cells infected with *L. monocytogenes* and *S. typhimurium* were washed once with PBS and given fresh DMEM medium containing 50 μg/ml gentamicin to kill the extracellular bacteria. The cells infected with *S. aureus* strains were washed once with PBS and given fresh DMEM medium containing 100 μg/ml gentamicin. At 4 hpi, cells were collected for RNA isolation. For macrophage infection with *F. novicida*, pBMDMs were infected with *F. novicida* at a MOI of 100. After 4 hpi, the cells were washed once with PBS and given fresh DMEM medium containing 50 μg/ml gentamicin, and the cells were collected at 6 hpi for RNA isolation.

For the study the inflammatory capacity or intracellular survive of *S*. aureus, either 1×10^6^ iBMDM or pBMDMs were plated in 6-well tissue culture plates and incubated at 37 °C in 5% carbon dioxide (CO_2_) overnight. Mid-exponential phase *S. aureus* cultures were washed once with sterile phosphate-buffered saline (PBS) and then re-suspended in DMEM medium. The macrophages were infected by *S. aureus* at a multiplicity of infection of 5 or 20. After a 1 hr incubation, the cells were washed twice with PBS and given fresh DMEM medium containing 100 μg/ml gentamicin to kill the extracellular bacteria. At various times post-infection, supernatants were collected to study macrophage cell death by lactate dehydrogenase (LDH) release assays (Promega, USA) ^66^, IFN-β protein expression by luciferase bioassays ^2^, and cells were collected for RNA isolation. For the luciferase bioassay, supernatants were applied in various dilutions to the IFN responsive ISRE-L929 cells (5×10^4^ cells/well, 96-well plate).The cells were lysed by TNT lysis and added to luciferase substrate solution and luminescence was measured using a Synergy HT (BioTek, USA).For intracellular CFU enumeration, macrophages were lysed in PBS containing 0.1% Triton X-100 and incubated at 37 °C for 5 min. Appropriate dilutions were plated on BHI agar plates and incubated at 37 °C overnight and CFU were enumerated.

### Laboratory *S. aureus* SCVs construction

*S. aureus* Newman Δ*thyA* and Δ*hemB* strains were constructed by allelic exchange using pIMAY^67^. Briefly, primers were designed to amplify upstream and downstream regions flanking the gene(s) of interest (either *thyA* or *hemB*) to delete the entire open reading frame (Supplemental Table 2). Genomic DNA from *S. aureus* strain Newman was used as a template to generate fragments consisting of linked sequences upstream and downstream of either *thyA* or *hemB* (Δ*thyA* or Δ*hemB* cassette) by overlapping PCR. The amplimer was cloned into pIMAY and transformed via electroporation into *E. coli* DH10B and then transformed into *S. aureus* strain RN4220. Transduction of *S. aureus* strain Newman was accomplished by first propagating phage ϕ-11 (to package the pIMAY vector) in RN4220 that had been successfully transformed with pIMAY_deletion cassette (Δ*thyA* or Δ*hemB*). Allelic exchange of either Δ*thyA* or Δ*hemB* was achieved by subsequent generalized phage transduction of the appropriate ϕ-11 pIMAY_deletion cassette vector into strain Newman. The resulting SCV strains were confirmed by PCR and further tested for auxotrophy for either thymidine or hemin; both showed attenuated growth in vitro compared with the wild-type Newman strain in the absence of either thymidine or hemin.

### *In vivo S. aureus* infection and bacterial burden estimation

Mice aged 8-12 weeks were used in infection experiments. Prior to animal experiments, overnight cultures of NC, Δ*thyA* and Δ*hemB* were back diluted twice (OD_600_=0.15) and grown for 2 h at 37 °C with 5 µg/mL of thymidine and 1 µg/mL hemin in BHI broth, respectively, to reach mid-exponential growth. Subsequently, the bacteria were washed once with PBS. Cultures were diluted in 30 μL PBS to achieve a final inoculum of 5×10^8^ bacteria and used to infect mice via the intranasal route while the animals were anesthetized with isoflurane. Mice were euthanized and lungs collected at the indicated time points. Lungs were homogenized and lysed in 5 mL PBS using a Tissue Tearor Homogenizer. Bacterial burdens were enumerated by plating dilutions on BHI plates containing thymidine or hemin to ensure the normal growth of SCVs.

### Repeat infection and histology

Mice aged 8-12 weeks were either intranasally instilled with 2.5×10^7^ bacteria in 30 μL PBS or 30 μL PBS every 4 days for a total of 5 infections. At day 16, mice were euthanized by CO_2_ exposure, the lungs were inflated with 1 mL cold 4% formalin and the trachea were tied up using silk string. Whole lungs were immersion fixed in 4% formalin, paraffin embedded, sectioned, stained with hematoxylin and eosin (H&E) and then evaluated for lung inflammation and injury. Lungs were scored semi-quantitatively in a blinded fashion for lung inflammation which includes perivascular/peribronchiolar inflammation, intrabronchiolar inflammation, interstitial inflammation, intraalveolar inflammation and severity/extent using a 1-4 scale in which 1 through 4 generally indicated “minimal”, “mild”, “moderate”, and “severe” changes, respectively. The lungs were also scored semi-quantitatively for lung injury which includes hemorrhage and collagen release. The representative images were scanned by Keyence BZ-X710 microscope and image brightness and contrast were adjusted using Photoshop applied to the entire image. Original magnification was as stated.

### TaqMan array experiments and relative quantitative real-time PCR

The left lungs of mice were sliced into pieces no wider than 0.5 cm and dropped into RNAlater (Thermo Fisher Scientific, USA) for RNA stabilization. At the time of RNA isolation, the samples were removed from RNAlater and homogenized by Bead Mill homogenizer (Thermo Fisher Scientific, USA) in Lysis buffer from NucleoSpin RNA Plus Kit (Clontech, USA). RNA from the lung homogenate or macrophages was isolated using NucleoSpin RNA Plus Kit (Clontech, USA) followed by cDNA synthesis using iScript cDNA synthesis Kits (Bio-Rad, USA) following manufacturer’s instructions. qRT-PCR was performed using TaqMan Gene expression master mix (Thermo Fisher Scientific, USA). Taqman probes for mouse genes were pre-designed assays from Thermo Fisher Scientific. To calculate mRNA fold change, transcript levels were determined by normalizing to mouse *Hprt* using the 2^(–ΔΔCt)^ method. Immune gene expression profiling was performed using TaqMan Array Micro Fluidic Cards (Applied Biosystems, USA) containing probe sets for 90 immune genes and 6 endogenous controls.

### Flowcytometry analysis for BAL neutrophils

Mice were euthanized at the indicated time by CO_2_ asphyxiation and lungs were lavaged with 0.5 ml ice-cold PBS twice. A total of 1 mL of lavage fluid was centrifuged at 2,000 g for 2 min. Collected cells were re-suspended in 100 µL Red Cell Lysis Buffer (BioLegend, USA) and incubated for 5 min. Cells lysis was stopped by adding 400 μL Cell Staining Buffer (BioLegend, USA). Cells were then centrifuge for 2 min at 2,000 g for 2 min and re-suspend in Cell Staining Buffer. Viable cells were counted using the Automatic cell counter (Bio-Rad) after trypan blue staining. Fc receptors were blocked by incubating cells with TruStain FcX™ PLUS anti-mouse CD16/32 antibody (Biolegend, USA) for 10 min. Cells were stained using the Zombie Green Fixable Viability Kit (BioLegend, USA) for 10 min to detect dead cells. Cells were washed twice with 500 µL PBS and resuspend in Cell Staining Buffer. Cells were thereafter stained with antibodies for the desired surface markers for 20 min on ice in the dark. Flow cytometric data were acquired on Cytek Aurora and analyzed using the FlowJo software. The following antibodies specific for the cell surface antigens from Biolegend were used for the flow cytometry: Brilliant Violet 421 anti-mouse Ly-6G, PE/Dazzle 594 anti-mouse/human CD11b, PerCP/Cyanine5.5 anti-mouse CD45, PE anti-mouse F4/80. 10, 000 live singlet cells were pre-gated. Neutrophils were identified as live, CD45^+^, F4/80^-^, CD11b^+^, Ly6G^+^ (Extended Data Fig. 5). The neutrophil numbers were calculated according to total cells in BALF and the percent of neutrophil identified by FACS assay.

### Intranasal c-di-AMP administration

C57BL/6J WT or *Sting*^-/-^ mice aged of 8-12 weeks were anesthetized under isoflurane. Mice were intranasally instilled with either PBS containing 15 μg c-di-AMP and 15 μg Pam3CSK4 or PBS containing 15 μg Pam3CSK4 and sacrificed at 8 hrs after treatment for BAL. The neutrophil numbers in BALF were enumerated by FACS assay and CXCL10, CCL5 and IL-6 concentrations in the BALF were measured by using the Mouse Quantikine ELISA Kit (R&D, USA).

### Neutrophil elastase assay

Elastase activity was measure by spectrofluorometrically monitoring the hydrolysis product of the fluorogenic substrate MeOSuc-AAPV-AMC (Cayman, USA). BALF was centrifuged at the maximum speed to remove the cell debris, then 20 µL BALF was mixed with 80 µL substrate solution (0.1 mM MeOSuc-AAPV-AMC, 0.5 M NaCl, 0.01 M CaCl_2_, 10% DMSO, 0.05 M Tris, pH 7.5) in white bottom plates. The kinetics of substrate cleavage (increase in fluorescence of the liberated 7-amino-4-methylcoumarin, AMC) was measured using a fluorometer (BioTek, USA) set at 370 nm excitation and 460 nm emission at 37°C. Porcine elastase (Sigma, USA) was used to generate a standard curve.

### Quantification of c-di-AMP

Overnight culture of NC, Δ*thyA* and Δ*hemB* strains grown in BHI supplemented with 5 μg/mL of thymidine were washed twice by PBS, then diluted into fresh BHI containing various concentration of thymidine (1.25, 2.5 and 5 μg/mL) with the initial OD_600_ of 0.05, and grown at 37 °C with shaking for 4 hrs. The bacteria were collected by centrifugation (2,700 × g, 5 min) and used for c-di-AMP extraction or macrophage infection. Macrophages infected with bacteria were collected by incubating in 0.05% Trypsin-EDTA for 5 min. Both macrophages and bacteria were re-suspended in methanol and lysed by sonication. Heavy-labeled (^3^H) c-di-AMP was added into the extract as the internal control. Bacterial lysates were centrifuged to collect cell-free supernatants. The quantification of c-di-AMP by LC-MS was performed as previously described^68^.

### Bioinformatic analyses

All numerical data were analyzed and plotted using GraphPad Prism 6.0 software. Statistical parameters are reported in the Figure Legends.

## Data availability

This study did not generate or analyze new datasets or codes.

## Acknowledgments

We would like to thank Daniel B. Stetson for providing *Tmem173*^-/-^ and *cGAS*^-/-^ mice. We thank Jessica M. Snyder at the Department of Comparative Medicine, University of Washington for histological scoring. We thank Samantha Hopp, Melissa Locke, Chelsea Stamm, Yaxi Wang for the help with either designing or performing experiments. Q.T. was supported by the University of Washington Cystic Fibrosis Foundation RDP Fellowship (SINGH15R0). This work was supported by a Pilot and Feasibility Award from the Cystic Fibrosis Foundation (WOODWA16I0) and NIH Grant R01 AI139071.

## Author contributions

Q.T. and J.J.W. conceived and designed the research. Q.T., M.R.P., M.K.T., and F.A.Q. performed experiments. Q.T., M.K.T., A.P.M. and J.J.W. provided key insights. D.W. and L.R.H provided key tools and reagents. Q.T. and J.J.W. analyzed the data and wrote the paper. All authors reviewed the manuscript prior to submission.

## Competing interests

The authors declare no competing interests.

## Notes

### Competing Interest Statement

The authors have declared no competing interest.

### Summary of Updates

We removed Figure 1-2 of the old version to supplemental data; we also deleted Figure 7 of the old version. We added 2 new figures (Figure 1 and Figure 2) to the new version, we also added more panels to the remaining figures (the current Figure 3, 5, 6). We also revised the abstract, introduction, result, and discussion sections.

## References

1. Anderson, R., Tintinger, G., Cockeran, R., Potjo, M. & Feldman, C. Beneficial and harmful interactions of antibiotics with microbial pathogens and the host innate immune system. Pharmaceuticals 3, 1694–1710 (2010).

2. Woodward, J. J., Lavarone, A. T. & Portnoy, D. A. C-di-AMP secreted by intracellular Listeria monocytogenes activates a host type I interferon response. Science (80). (2010) doi:10.1126/science.1189801.

3. Burdette, D. L. et al. STING is a direct innate immune sensor of cyclic di-GMP. Nature (2011) doi:10.1038/nature10429.

4. Ishikawa, H. & Barber, G. N. STING is an endoplasmic reticulum adaptor that facilitates innate immune signalling. Nature (2008) doi:10.1038/nature07317.

5. Sun, L., Wu, J., Du, F., Chen, X. & Chen, Z. J. Cyclic GMP-AMP synthase is a cytosolic DNA sensor that activates the type I interferon pathway. Science (80). (2013) doi:10.1126/science.1232458.

6. Wu, J. et al. Cyclic GMP-AMP is an endogenous second messenger in innate immune signaling by cytosolic DNA. Science (80-.). (2013) doi:10.1126/science.1229963.

7. Zhang, X. et al. Cyclic GMP-AMP containing mixed Phosphodiester linkages is an endogenous high-affinity ligand for STING. Mol. Cell (2013) doi:10.1016/j.molcel.2013.05.022.

8. Gaidt, M. M. et al. The DNA inflammasome in human myeloid cells is initiated by a sting-cell death program upstream of NLRP3. Cell (2017) doi:10.1016/j.cell.2017.09.039.

9. Gui, X. et al. Autophagy induction via STING trafficking is a primordial function of the cGAS pathway. Nature (2019) doi:10.1038/s41586-019-1006-9.

10. Balka, K. R. et al. TBK1 and IKKε Act redundantly to mediate STING-induced NF-κB responses in myeloid cells. Cell Rep. (2020) doi:10.1016/j.celrep.2020.03.056.

11. Chen, H. et al. Activation of STAT6 by STING is critical for antiviral innate immunity. Cell (2011) doi:10.1016/j.cell.2011.09.022.

12. Gulen, M. F. et al. Signalling strength determines proapoptotic functions of STING. Nat. Commun. 8, 427 (2017).

13. Cheng, Z. et al. The interactions between cGAS-STING pathway and pathogens. Signal Transduct. Target. Ther. 5, 91 (2020).

14. Ma, Z. & Damania, B. The cGAS-STING defense pathway and its counteraction by viruses. Cell Host Microbe 19, 150–158 (2016).

15. Sun, Y. & Cheng, Y. STING or Sting: cGAS-STING-mediated immune response to protozoan parasites. Trends Parasitol. 36, 773–784 (2020).

16. Frémond, M.-L. et al. Overview of STING-associated vasculopathy with onset in infancy (SAVI) among 21 patients. J. Allergy Clin. Immunol. Pract. 9, 803–818.e11 (2021).

17. Liu, Y. et al. Activated STING in a vascular and pulmonary syndrome. N. Engl. J. Med. (2014) doi:10.1056/NEJMoa1312625.

18. Gao, D. et al. Activation of cyclic GMP-AMP synthase by self-DNA causes autoimmune diseases. Proc. Natl. Acad. Sci. 112, E5699--E5705 (2015).

19. Stülke, J. & Krüger, L. Cyclic di-AMP signaling in bacteria. Annu. Rev. Microbiol. 74, 159–179 (2020).

20. Yin, W. et al. Cyclic di-AMP signaling in bacteria. FEMS Microbiol. Rev. 44, 701–724 (2020).

21. Ba, X. et al. Truncation of GdpP mediates β-lactam resistance in clinical isolates of Staphylococcus aureus. J. Antimicrob. Chemother. 74, 1182–1191 (2019).

22. Sommer, A. et al. Mutations in the gdpP gene are a clinically relevant mechanism for β-lactam resistance in meticillin-resistant Staphylococcus aureus lacking mec determinants. Microb. genomics 7, (2021).

23. Argudín, M. A. et al. Genetic diversity among Staphylococcus aureus isolates showing oxacillin and/or cefoxitin resistance not linked to the presence of mec Genes. Antimicrob. Agents Chemother. 62, e00091–18 (2018).

24. Sun, D., Jeannot, K., Xiao, Y. & Knapp, C. W. Editorial: Horizontal gene transfer mediated bacterial antibiotic resistance. Front. Microbiol. 10, 1933 (2019).

25. Proctor, R. A. et al. Small colony variants: A pathogenic form of bacteria that facilitates persistent and recurrent infections. Nature Reviews Microbiology (2006) doi:10.1038/nrmicro1384.

26. Goerke, C. & Wolz, C. Adaptation of Staphylococcus aureus to the cystic fibrosis lung. International Journal of Medical Microbiology (2010) doi:10.1016/j.ijmm.2010.08.003.

27. Wolter, D. J. et al. Prevalence and clinical associations of Staphylococcus aureus small-colony variant respiratory infection in children with cystic fibrosis (SCVSA): a multicentre, observational study. Lancet Respir. Med. 7, 1027–1038 (2019).

28. Wolter, D. J. et al. Staphylococcus aureus small-colony variants are independently associated with worse lung disease in children with cystic fibrosis. Clin. Infect. Dis. (2013) doi:10.1093/cid/cit270.

29. Hawser, S., Lociuro, S. & Islam, K. Dihydrofolate reductase inhibitors as antibacterial agents. Biochem. Pharmacol. 71, 941–948 (2006).

30. Fernández-Villa, D., Aguilar, M. R. & Rojo, L. Folic Acid Antagonists: Antimicrobial and Immunomodulating Mechanisms and Applications. Int. J. Mol. Sci. 20, (2019).

31. Kompis, I. M., Islam, K. & Then, R. L. DNA and RNA Synthesis: Antifolates. Chem. Rev. 105, 593–620 (2005).

32. Besier, S. et al. Thymidine-dependent Staphylococcus aureus small-colony variants: human pathogens that are relevant not only in cases of cystic fibrosis lung disease. J. Clin. Microbiol. 46, 3829–3832 (2008).

33. Kriegeskorte, A. et al. Thymidine-dependent Staphylococcus aureus small-colony variants are induced by trimethoprim-sulfamethoxazole (SXT) and have increased fitness during SXT challenge. Antimicrob. Agents Chemother. (2015) doi:10.1128/AAC.00742-15.

34. Besier, S. et al. Prevalence and Clinical Significance of Staphylococcus aureus small-colony variants in cystic fibrosis lung disease. J. Clin. Microbiol. 45, 168–172 (2007).

35. Precit, M. R. et al. Optimized in vitro antibiotic susceptibility testing method for small-colony variant Staphylococcus aureus. Antimicrob. Agents Chemother. (2016) doi:10.1128/AAC.02330-15.

36. Davies, B. W., Bogard, R. W., Young, T. S. & Mekalanos, J. J. Coordinated regulation of accessory genetic elements produces cyclic di-nucleotides for V. cholerae virulence. Cell (2012) doi:10.1016/j.cell.2012.01.053.

37. Whiteley, A. T. et al. Bacterial cGAS-like enzymes synthesize diverse nucleotide signals. Nature (2019) doi:10.1038/s41586-019-0953-5.

38. Corrigan, R. M., Abbott, J. C., Burhenne, H., Kaever, V. & Gründling, A. C-di-AMP is a new second messenger in Staphylococcus aureus with a role in controlling cell size and envelope stress. PLoS Pathog. (2011) doi:10.1371/journal.ppat.1002217.

39. Holland, L. M. et al. A staphylococcal GGDEF domain protein regulates biofilm formation independently of cyclic dimeric GMP. J. Bacteriol. (2008) doi:10.1128/JB.00375-08.

40. Witte, C. E. et al. Cyclic di-AMP is critical for Listeria monocytogenes growth, cell wall homeostasis, and establishment of infection. MBio (2013) doi:10.1128/mBio.00282-13.

41. Moretti, J. et al. STING senses microbial viability to orchestrate stress-mediated autophagy of the endoplasmic reticulum. Cell (2017) doi:10.1016/j.cell.2017.09.034.

42. McFarland, A. P. et al. Sensing of bacterial cyclic dinucleotides by the oxidoreductase RECON promotes NF-κB activation and shapes a proinflammatory antibacterial state. Immunity (2017) doi:10.1016/j.immuni.2017.02.014.

43. Zander, J. et al. Influence of dTMP on the phenotypic appearance and intracellular persistence of Staphylococcus aureus. Infect. Immun. (2008) doi:10.1128/IAI.01075-07.

44. Kriegeskorte, A. et al. Inactivation of thyA in Staphylococcus aureus attenuates virulence and has a strong impact on metabolism and virulence gene expression. MBio (2014) doi:10.1128/mBio.01447-14.

45. Abe, T. & Barber, G. N. Cytosolic-DNA-mediated, STING-dependent proinflammatory gene induction necessitates canonical NF-κB Activation through TBK1. J. Virol. (2014) doi:10.1128/jvi.00037-14.

46. Yum, S., Li, M., Fang, Y. & Chen, Z. J. TBK1 recruitment to STING activates both IRF3 and NF-κB that mediate immune defense against tumors and viral infections. Proc. Natl. Acad. Sci. 118, (2021).

47. McGuinness, W. A., Kobayashi, S. D. & DeLeo, F. R. Evasion of neutrophil killing by Staphylococcus aureus. Pathogens (2016) doi:10.3390/pathogens5010032.

48. Kaplanski, G., Marin, V., Montero-Julian, F., Mantovani, A. & Farnarier, C. IL-6: a regulator of the transition from neutrophil to monocyte recruitment during inflammation. Trends Immunol. 24, 25–29 (2003).

49. Pan, Z. Z., Parkyn, L., Ray, A. & Ray, P. Inducible lung-specific expression of RANTES: preferential recruitment of neutrophils. Am. J. Physiol. Lung Cell. Mol. Physiol. 279, L658–66 (2000).

50. Ichikawa, A. et al. CXCL10-CXCR3 enhances the development of neutrophil-mediated fulminant lung injury of viral and nonviral origin. Am. J. Respir. Crit. Care Med. 187, 65–77 (2013).

51. Dengler, V. et al. Mutation in the c-di-AMP cyclase DacA affects fitness and resistance of methicillin resistant Staphylococcus aureus. PLoS One (2013) doi:10.1371/journal.pone.0073512.

52. Besier, S. et al. Hypermutability and antibiotic resistance in thymidinedependent small-colony-variants of Staphylococcus aureus. Int. J. Med. Microbiol. (2009).

53. Oppenheimer-Shaanan, Y., Wexselblatt, E., Katzhendler, J., Yavin, E. & Ben-Yehuda, S. C-di-AMP reports DNA integrity during sporulation in Bacillus subtilis. EMBO Rep. (2011) doi:10.1038/embor.2011.77.

54. Zeden, M. S. et al. Cyclic di-adenosine monophosphate (c-di-AMP) is required for osmotic regulation in Staphylococcus aureus but dispensable for viability in anaerobic conditions. J. Biol. Chem. (2018) doi:10.1074/jbc.M117.818716.

55. Choi, P. H., Sureka, K., Woodward, J. J. & Tong, L. Molecular basis for the recognition of cyclic-di-AMP by PstA, a PII-like signal transduction protein. Microbiologyopen (2015) doi:10.1002/mbo3.243.

56. Müller, M., Hopfner, K. P. & Witte, G. C-di-AMP recognition by Staphylococcus aureus PstA. FEBS Lett. (2015) doi:10.1016/j.febslet.2014.11.022.

57. Orman, K. L. & English, B. K. Effects of antibiotic class on the macrophage inflammatory response to Streptococcus pneumoniae. J. Infect. Dis. 182, 1561–1565 (2000).

58. Wolf, A. J., Liu, G. Y. & Underhill, D. M. Inflammatory properties of antibiotic-treated bacteria. J. Leukoc. Biol. 101, 127–134 (2017).

59. Porto, B. N. & Stein, R. T. Neutrophil extracellular traps in pulmonary diseases: Too much of a good thing? Front. Immunol. (2016) doi:10.3389/fimmu.2016.00311.

60. Phillipson, M. & Kubes, P. The neutrophil in vascular inflammation. Nat. Med. 17, 1381–1390 (2011).

61. Askarian, F., Wagner, T., Johannessen, M. & Nizet, V. Staphylococcus aureus modulation of innate immune responses through Toll-like (TLR), (NOD)-like (NLR) and C-type lectin (CLR) receptors. FEMS Microbiol. Rev. 42, 656–671 (2018).

62. Brandt, S. L., Putnam, N. E., Cassat, J. E. & Serezani, C. H. Innate immunity to Staphylococcus aureus: evolving paradigms in soft tissue and invasive infections. J. Immunol. 200, 3871LP–3880 (2018).

63. Kahl, B. C. et al. Thymidine-dependent small-colony variants of Staphylococcus aureus exhibit gross morphological and ultrastructural changes consistent with impaired cell separation. J. Clin. Microbiol. (2003) doi:10.1128/JCM.41.1.410-413.2003.

64. Weischenfeldt, J. & Porse, B. Bone marrow-derived macrophages (BMM): Isolation and applications. Cold Spring Harb. Protoc. (2008) doi:10.1101/pdb.prot5080.

65. Brunette, R. L. et al. Extensive evolutionary and functional diversity among mammalian AIM2-like receptors. J. Exp. Med. (2012) doi:10.1084/jem.20121960.

66. Smith, S. M., Wunder, M. B., Norris, D. A. & Shellman, Y. G. A simple protocol for using a LDH-Based cytotoxicity assay to assess the effects of death and growth inhibition at the same time. PLoS One (2011) doi:10.1371/journal.pone.0026908.

67. Monk, I. R., Shah, I. M., Xu, M., Tan, M. W. & Foster, T. J. Transforming the untransformable: Application of direct transformation to manipulate genetically Staphylococcus aureus and Staphylococcus epidermidis. MBio (2012) doi:10.1128/mBio.00277-11.

68. Huynh, T. A. N. et al. An HD-domain phosphodiesterase mediates cooperative hydrolysis of c-di-AMP to affect bacterial growth and virulence. Proc. Natl. Acad. Sci. U. S. A. (2015) doi:10.1073/pnas.1416485112.

