## Supplemental figures and Table for "Thymidine starvation promotes c-di-AMP dependent inflammation during infection"

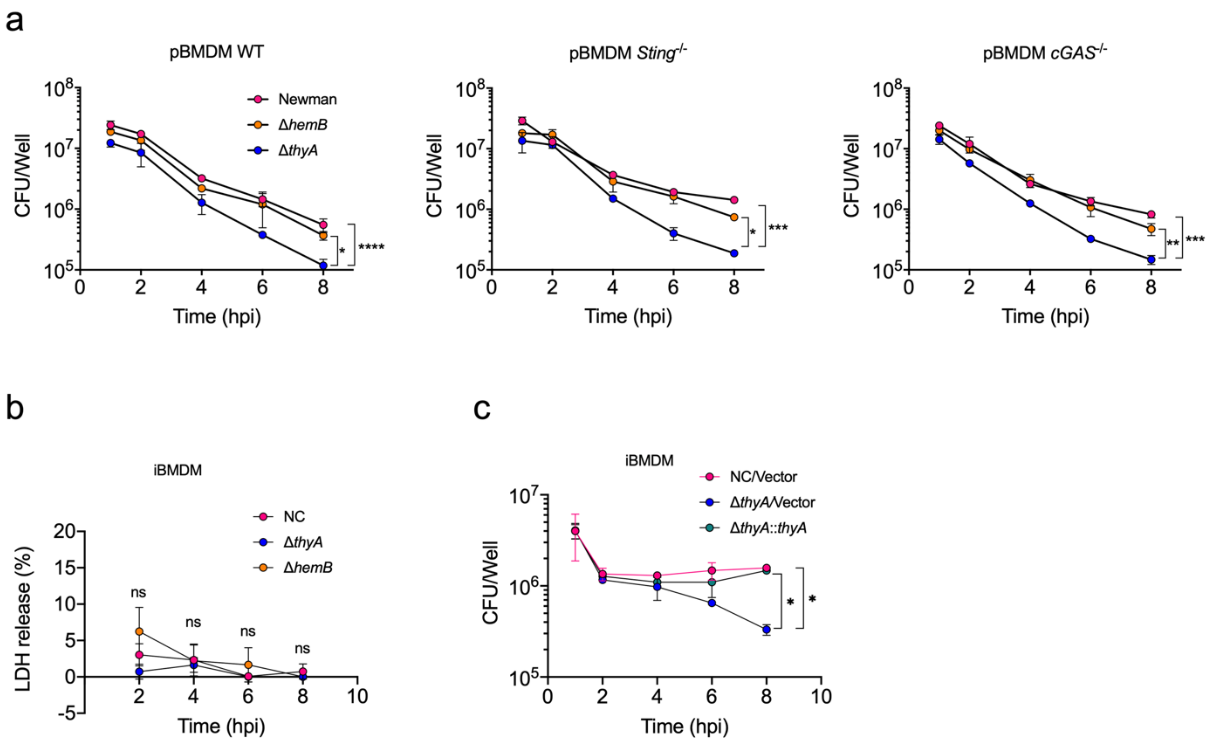


**Extended Data Fig. 1: Δ*thyA* exhibits decreased survival during macrophage infection**.

**a**, Bacterial survival of NC, Δ*thyA*, and Δ*hemB S. aureus* Newman strains in in WT, *Sting*$^{-/-}$ and *cGas*$^{-/-}$ pBMDMs. **b**, The iBMDM were infected by NC, Δ*thyA*, and Δ*hemB S. aureus* strains at a multiplicity of infection (MOI) of 5. LDH release of iBMDMs infected with indicated *S. aureus* Newman strains normalized to 100% lysis controls. **c**, Bacterial survival curves of NC, Δ*thyA*, and Δ*thyA::thyA S. aureus* Newman strains in iBMDM cells. The iBMDM were infected by *S. aureus* at a MOI of 5. For all panels, mean values of duplicate are plotted and error bars indicate ±SD. *P* values were calculated using Two-way ANOVA analysis. Asterisks indicate that differences are statistically significant (*, $P < 0.05$; **, $P < 0.01$; ***, $P < 0.001$, ****, $P < 0.0001$), and ns indicates no significant difference.

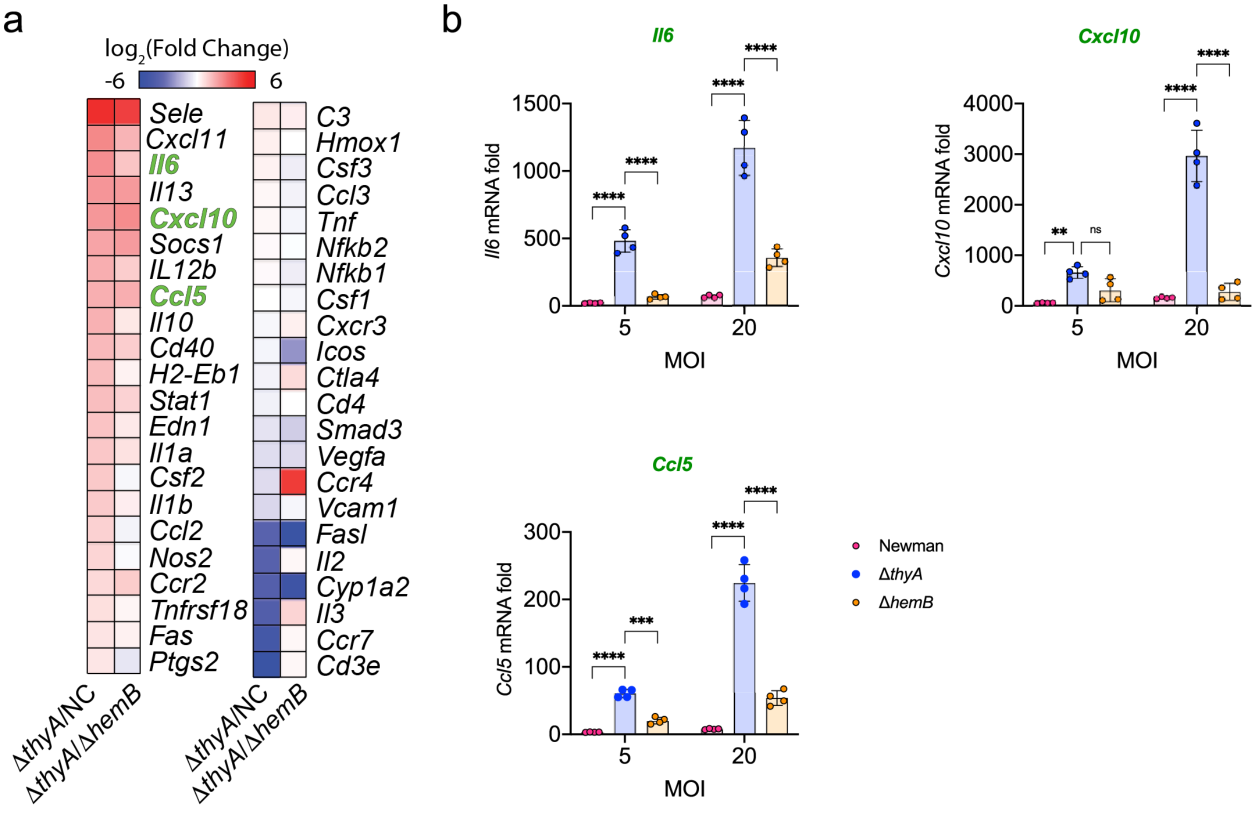


**Extended Data Fig. 2: Δ*thyA* promotes elevated inflammation during macrophage infection.**

**a**, Heatmap of Taqman gene expression analysis of WT iBMDMs infected with NC, Δ*thyA*, and Δ*hemB* *S. aureus* Newman strains. Values are the $\log_2$ ratio of differences in the transcription of the indicated genes (Δ*thyA* vs NC and Δ*thyA* vs Δ*hemB*). Heatmaps were generated using Morpheus (https://software.broadinstitute.org/morpheus). **b**, Transcriptional differences for genes labeled green in panel **a**) were confirmed by qRT-PCR in iBMDMs with *S. aureus* infection at MOI of either 5 or 20. For all panels, error bars indicate ±SD, $P$ values were calculated using Two-way ANOVA analysis. Asterisks indicate that differences are statistically significant ( **, $P < 0.01$; ***, $P < 0.001$, ****, $P < 0.0001$), and ns indicates no significant difference.

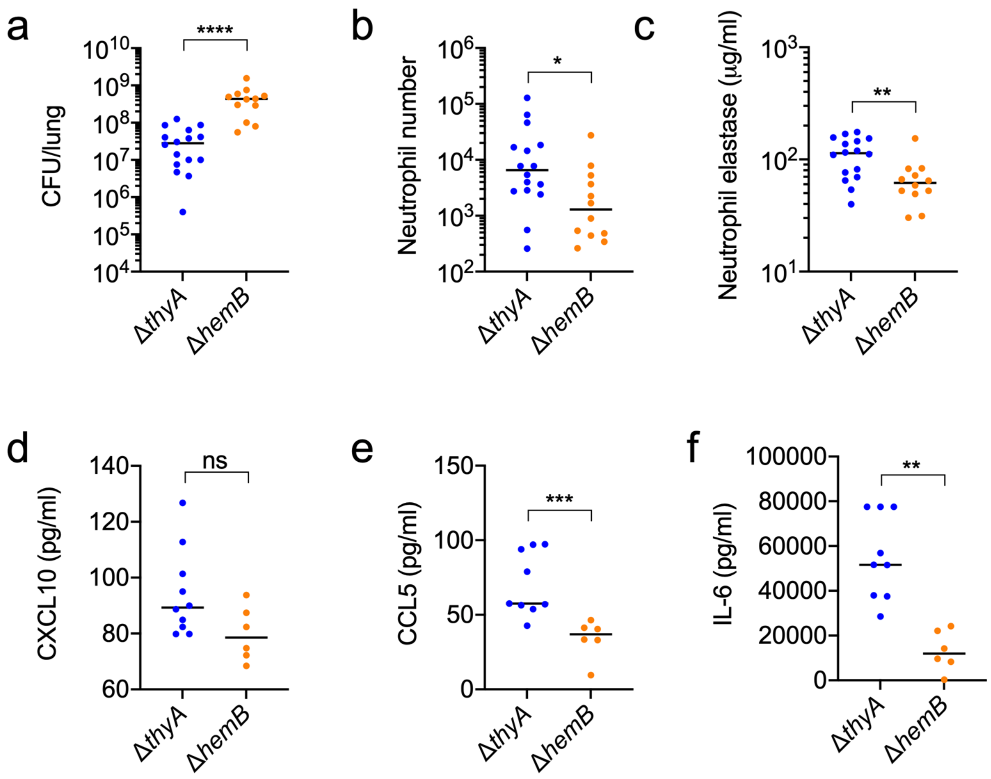


**Extended Data Fig. 3: Δ*thyA* induces higher airway neutrophil infiltration than Δ*hemB*. a**, WT mice were intranasally infected with the Δ*thyA* or Δ*hemB* strains. The CFU recovery from the right lung were enumerated at 8 hpi. **b**, The neutrophils in BALF of the mice infected as in panel (**a**) were identified and enumerated by FACS assay. **c**, Active neutrophil elastase concentration in BALF form mice infected as in panel (**a**). **d-f**, BALF CXCL10, CCL5 and IL-6 concentration of mice infected as panel (**a**) determined by ELISA. For all panels, the horizontal black bar is the median of the data. *P* values were calculated using Mann-Whitney analysis. Asterisks indicate that differences are statistically significant (**, $P < 0.01$).

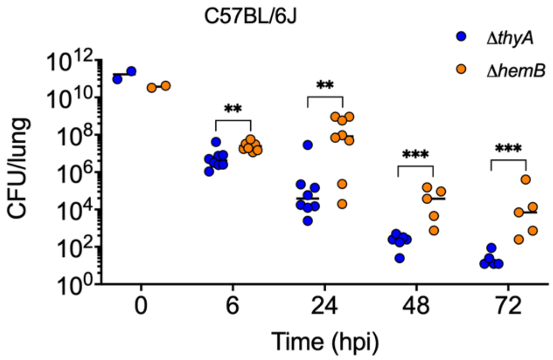


**Extended Data Fig. 4: Δ*thyA* exhibits decreased survival during murine lung infection.**
Bacterial recovery from the lung of WT C57BL/6J mice intranasally infected with Δ*thyA* and Δ*hemB*. Horizontal black bars are the median of the measured bacterial burdens. *P* values were calculated using Mann-Whitney analysis. Asterisks indicate that differences are statistically significant (**, $P < 0.01$; ***, $P < 0.001$), and ns indicates no significant difference.

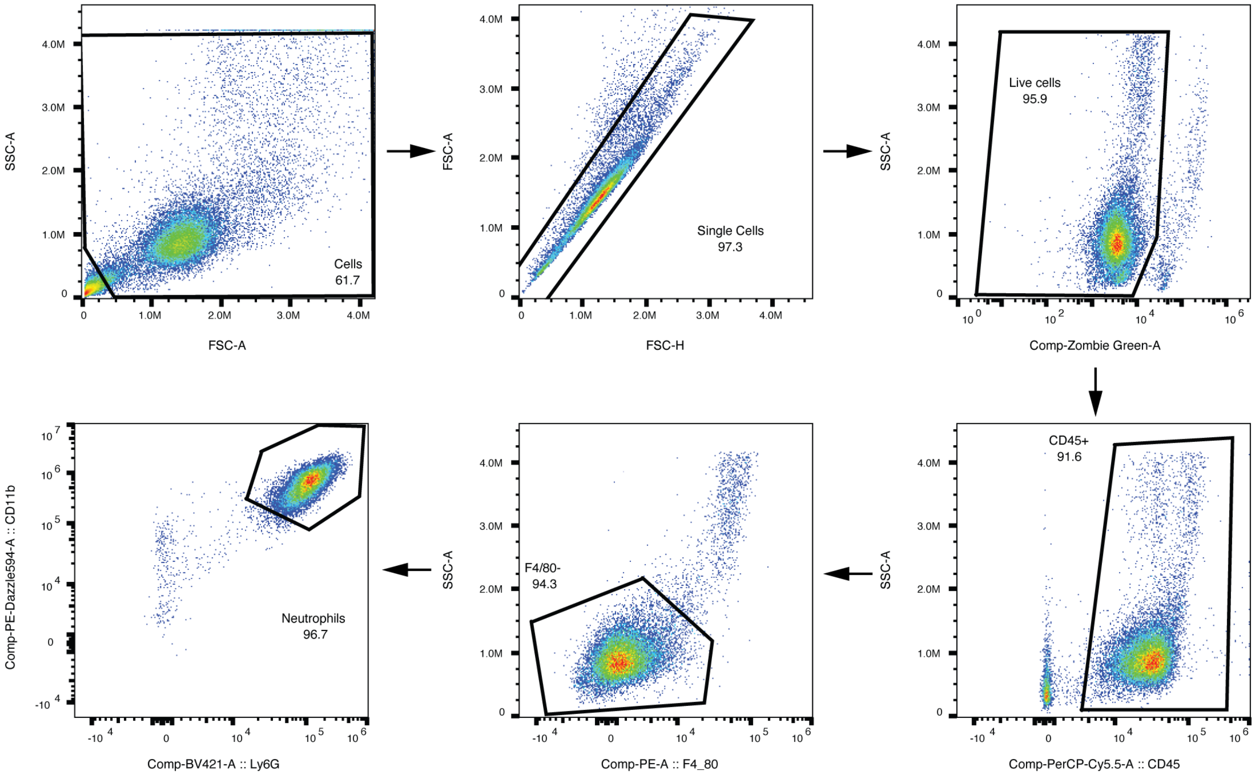


**Extended Data Fig. 5: Flow cytometry gating strategy.**
Neutrophils were identified as live, CD45+, F4/80-, CD11b+, Ly6G+.

**Table S1 Histological scores**

| Slide ID | Inflammation‡ | | | | Overall severity | Extent | Severity[e] × Extent[f] | Total inflammation | NOTES |
|---|---|---|---|---|---|---|---|---|---|
| | **Perivascular/ Peribronchiolar** | **Intrabronchiolar** | **Interalveolar** | **Interstitial** | | | | | |
| PBS_1 | 1 | 1 | 1 | 1 | 1 | 1 | 1 | 5 | Mild, fibrous nodules |
| PBS_2 | 0 | 0 | 0 | 0 | 0 | 0 | 0 | 0 | Basically looks normal |
| PBS_3 | 1 | 0 | 1 | 1 | 1 | 1 | 1 | 4 | mild focal |
| PBS_4 | 0 | 1 | 0 | 0 | 0 | 0 | 0 | 1 | Focal minimal inflammation in one bronchiole with refractile material |
| PBS_5 | 0 | 0 | 0 | 0 | 0 | 0 | 0 | 0 | Normal |
| NC_1 | 2 | 0 | 2 | 1 | 2 | 2 | 4 | 9 | Mild |
| NC_2 | 2 | 0 | 1 | 1 | 1 | 1 | 1 | 5 | Mild |
| NC_3 | 3 | 1 | 3 | 2 | 2 | 3 | 6 | 15 | Moderate |
| NC_4 | 3 | 1 | 2 | 2 | 2 | 2 | 4 | 12 | Moderate |
| NC_5 | 3 | 1 | 2 | 2 | 2 | 3 | 6 | 14 | Moderate |
| NC_6 | 2 | 0 | 2 | 2 | 2 | 2 | 4 | 10 | Rare fibrous nodules |
| D*thyA*_1 | 2 | 1 | 2 | 3 | 2 | 3 | 6 | 14 | |
| D*thyA*_2 | 3 | 1 | 2 | 2 | 2 | 2 | 4 | 12 | |
| D*thyA*_3 | 2 | 3 | 3 | 2 | 2 | 3 | 6 | 16 | More intrabronchiolar inflammation |
| D*thyA*_4 | 2 | 3 | 2 | 2 | 2 | 3 | 6 | 15 | More intrabronchioalr inflammation |
| DthyA_5 | 3 | 1 | 2 | 2 | 2 | 3 | 6 | 14 | |
| D*thyA*_6 | 2 | 3 | 2 | 3 | 2 | 3 | 6 | 16 | More intrabronchiolar inflammation |
| D*hemB*_1 | 3 | 1 | 2 | 1 | 2 | 3 | 4 | 11 | |

| | | | | | | | | | |
|---|---|---|---|---|---|---|---|---|---|
| D*hemB*_2 | 2 | 0 | 2 | 2 | 2 | 2 | 4 | 10 | Mild |
| D*hemB*_3 | 3 | 2 | 3 | 3 | 2 | 3 | 6 | 17 | |
| D*hemB*_4 | 3 | 1 | 2 | 2 | 2 | 2 | 4 | 12 | |
| D*hemB*_5 | 2 | 1 | 2 | 2 | 2 | 2 | 4 | 11 | |
| D*hemB*_6 | 2 | 1 | 2 | 2 | 2 | 2 | 4 | 11 | |

‡

Inflammation:

| | Perivascular /Peribronchiolar | Intrabronchiolar | Interstitial | Intraalveolar |
|---|---|---|---|---|
| 1 | rare scattered cuffs of cells | rare scattered cells or focus of cells in 1 bronchiolar lumen | minimal | rare, scattered |
| 2 | cuffs of cells up to 15% | foci of cells expanding 2-3 bronchiolar lumens | mild thickening | affecting up to 15% |
| 3 | cuffs 15-33% with pv red blood cells | 4-10 bronchiolar lumens expanded by cells | moderate, involving increased interstitium around vessels | mf to c, expanding alveoli |
| 4 | >33% | severe, >10 | severe, >33% | severe, >33% |

*Severity: 0 = normal; 1 = few inflammatory cells (<5%); 2 = larger foci of inflammatory cells, with preservation of underlying architecture (5-15%); 3 = larger foci of inflammatory cells with mild changes of underlying vessel / alveolar wall or bronchiolar epithelium (includes perivascular, interstitial, or alveolar hemorrhage, and 16-33% affected); 4 = larger foci of inflammatory cells with disruption or loss of underlying architecture including necrosis and hemorrhage.

†Extent: 0 = none; 1 = rare, scattered <5%; 2 = larger focus or multifocal small foci 5 - 15%; 3 = 15-33%; 4 > 34%.

**Table S2. Oligonucleotides used in this paper**

| Primers | Sequences | Notes |
|---|---|---|
| **Allelic exchange vector construction** | | |
| *thyA* SOE FwdA | ATATGGTACCGAAGCAGTATCGGAGTATATG | KpnI |
| *thyA* SOE RevB | CTATGCAATGACTACATATGCGATAACACCTCATTTTC | |
| *thyA* SOE FwdC | GAGGTGTTATCGCATATGTAGTCATTGCATAGTTAGCTAAC | |
| *thyA* SOE RevD | ATATGCGGCCGCTATGGCTGGCTGACTTGTC | NotI |
| *hemB* SOE FwdA | ATATGGTACCGAAGCCAGTTGTAGTTATGAC | KpnI |
| *hemB* SOE RevB | CATAAATATAAAACCTTACATTTTTAGCCCCTACTTTC | |
| *hemB* SOE FwdC | GTAGGGGCTAAAAATGTAAGGTTTTATATTTATGATTTTCC | |
| *hemB* SOE RevD | ATATGCGGCCGCCATATTACCAGCAACAGGTTCTAC | NotI |
| ***S. aureus* mutant verification** | | |
| *thyA*-Fwd1 | GTATGGAAACAAATGGCAAACAGAAC | |
| *thyA*-Rev1 | GTACCACTTAATCCTGAAGAAAGATG | |
| *hemB*-Fwd1 | CAGAAATGGATGGTAGTTGTCAG | |
| *hemB*-Rev1 | GCACAATGATAACCGACTCTG | |
| **pEPSA5-*Lmpde* vector construction** | | |
| Lmpde-F | CGGAATTCAACGACTGGAAGGAGTTTTAATTAATGCCGATGGGAATACTGCTGTA | KpnI |
| Lmpde-R | GGGGTACCTTAgtgatgatgatgatgatgTGTTTCTCCCTTCCAATACG | EcoRI |
| **pBAV1K-E*-*dacA* vector construction** | | |
| *dacA-SOE-a* | GAAGCTAATTATAACAAGACGAACTCC | |
| *dacA-SOE-b* | CCTGCTAAGGAGGCAACAAGATGGATTTTTCCAACTTTTTTCAAAACC | |
| *dacA-SOE-c* | GGTTTTGAAAAAAGTTGGAAAAATCCATCTTGTTGCCTCCTTAGCAGG | |
| *dacA-SOE-d* | GAGGAGACTAGTTTATTTCACACCTTTCTTTTGAAAGC | |
| ***Tmem173**$^{-/-}$ **mouse genotyping** | | |
| STING common S1 | CAA TGC TCT CAT AGC CTT CAC TAT C | |
| STING WT AS1 | AGA ACG GAC AGC CAG TAA GTA TAC AG | |
| STING mutant AS1 | AAC TTC CTG ACT AGG GGA GGA GTA G | |
